# Developmental origin and local signals cooperate to determine septal astrocyte identity

**DOI:** 10.1101/2023.10.08.561428

**Authors:** Yajun Xie, Christopher M. Reid, Alejandro A. Granados, Miguel Turrero Garcia, Fiona Dale-Huang, Sarah M. Hanson, Walter Mancia, Jonathan Liu, Manal Adam, Olivia Mosto, Angela O. Pisco, Arturo Alvarez-Buylla, Corey C. Harwell

**Affiliations:** Department of Neurology, University of California, San Francisco, CA; Department of Neurological Surgery, University of California, San Francisco, CA; Eli and Edythe Broad Center of Regeneration Medicine and Stem Cell Research, San Francisco, CA; Department of Neurobiology, Harvard Medical School, Boston, MA; Ph.D. Program in Neuroscience, Harvard University, Boston, MA; Chan Zuckerberg Biohub San Francisco, San Francisco, CA

## Abstract

Astrocyte specification during development is influenced by both intrinsic and extrinsic factors, but the precise contribution of each remains poorly understood. Here we show that septal astrocytes from Nkx2.1 and Zic4 expressing progenitor zones are allocated into non-overlapping domains of the medial (MS) and lateral septal nuclei (LS) respectively. Astrocytes in these areas exhibit distinctive molecular and morphological features tailored to the unique cellular and synaptic circuit environment of each nucleus. Using single-nucleus (sn) RNA sequencing, we trace the developmental trajectories of cells in the septum and find that neurons and astrocytes undergo region and developmental stage-specific local cell-cell interactions. We show that expression of the classic morphogens Sonic hedgehog (Shh) and Fibroblast growth factors (Fgfs) by MS and LS neurons respectively, functions to promote the molecular specification of local astrocytes in each region. Finally, using heterotopic cell transplantation, we show that both morphological and molecular specifications of septal astrocytes are highly dependent on the local microenvironment, regardless of developmental origins. Our data highlights the complex interplay between intrinsic and extrinsic factors shaping astrocyte identities and illustrates the importance of the local environment in determining astrocyte functional specialization.

## INTRODUCTION

The septal area, situated in the ventral forebrain, governs a diverse array of behaviors related to memory, movement, mood and motivational processes (Kesner et al., 2021; Menon, Suss, Oliveira, Neumann, & Bludau, 2022; Mocellin & Mikulovic, 2021; Wirtshafter & Wilson, 2021). These multifaceted functions of the septum are thought to be mediated by diverse septal neuron cell types, which establish circuit connections with a wide range of brain regions, most notably the hippocampus and hypothalamic area nuclei (Muller & Remy, 2018; Sun et al., 2014; D. Wang et al., 2023). Over the past several years, there has been significant progress in delineating the roles of genetically defined septal neuron types (Magno et al., 2022; Turrero Garcia et al., 2021). However, there remains a significant gap in understanding the extent and functional relevance of glial cell diversity in the septum. It is now appreciated that glial cells significantly contribute to neural circuit function and behavior outputs (Hirrlinger & Nimmerjahn, 2022; Lawal, Ulloa Severino, & Eroglu, 2022; Lyon & Allen, 2021; Perez-Catalan, Doe, & Ackerman, 2021). Therefore, it is important to understand the heterogeneity of glial cell types within the septal area, to complement our growing knowledge of neuron cell type diversity. In this study, we generate a comprehensive catalog of septal cell types during postnatal development, advancing our understanding of the molecular diversity and cell type specification in this brain region.

Astrocytes are an abundant glial cell type extensively distributed throughout the central nervous system (CNS). There is a growing recognition of the morphological and molecular diversity exhibited by astrocytes in different CNS regions (Batiuk et al., 2020; Bayraktar, Fuentealba, Alvarez-Buylla, & Rowitch, 2014; Farmer & Murai, 2017; M. G. Holt, 2023; Miller, 2018). However, the developmental mechanisms underlying the diversification and functional specialization of astrocytes to support a wide range of neural circuits are not understood. It is widely recognized that astrocyte specification is shaped by the interplay between intrinsic factors related to developmental origins/progenitor lineages and extrinsic factors derived from the local cellular environment (Farmer et al., 2016; Markey, Saunders, Smuts, von Reyn, & Garcia, 2023; Zarei-Kheirabadi, Vaccaro, Rahimi-Movaghar, Kiani, & Baharvand, 2020). However, the precise roles and contributions of each of these factors have yet to be determined. By combining genetic fate-mapping with snRNA sequencing, we identified two astrocyte populations residing in the MS and LS respectively. These two astrocyte populations are distinguished by their anatomical allocation within the septum, their developmental origins, and their molecular and morphological properties. We traced the molecular specification of these two astrocyte populations over postnatal development and identified sets of functional genes that characterize each type. We found MS and LS astrocytes exhibit unique developmental trajectories and exclusive interactions with local neighboring neurons. Many of the genes that distinguish MS and LS astrocytes were downstream transcriptional targets of Shh and Fgf signaling, two neuron-derived secreted factors expressed in the MS and LS respectively. To determine the precise role of local environmental cues in specifying the distinctive features of MS and LS astrocytes, we performed heterotopic transplantations of cells derived from either lineage and found that transplanted astrocytes adopted morphological and molecular features of astrocytes in the host region, regardless of their developmental origins. Together, our findings indicate that local neuron-derived factors have a profound impact on shaping the specialized properties of astrocytes within their local environment and exemplify the remarkable plasticity of astrocytes to adapt to their surrounding microenvironment.

## RESULTS

### Nkx2.1 and Zic4-derived astrocytes occupy the MS and LS respectively

Nkx2.1+ progenitors are known to significantly contribute to the production of glial populations in ventral telencephalic regions (Minocha et al., 2017; Tsai et al., 2012). To determine the specific contribution of Nxk2.1 progenitors to septal astrocyte diversity, we used the Nkx2.1 Cre mouse line crossed with the Sun1GFP reporter line to fluorescently label cells with a history of Nkx2.1 expression. Immunostaining postnatal day (P) 30 Nkx2.1-Cre; Sun1GFP mice with the astrocyte marker Sox9 revealed that the majority of Sox9+ astrocytes (74%) in the MS have a history of Nkx2.1 expression (Turrero Garcia et al., 2021; Wei et al., 2012). In contrast, anatomical subdivisions of the LS have fewer Nkx2.1-derived astrocytes (dLS: 0.1%; iLS: 2%; vLS: 12%) (**Figure 1A and 1B**), mirroring oligodendrocyte lineage patterns (**Figures S1A and S1B**). Sun1GFP-positive astrocytes in the MS make up 18% of all Nkx2.1-derived cells (**Figure 1C**), consistent with Nkx2.1 progenitor contribution to other neuronal and glial cell types within the MS (Magno et al., 2017; Turrero Garcia et al., 2021) (**Figure S1C**). Zic-expressing progenitors produce the majority of neurons in the LS (Inoue, Ota, Ogawa, Mikoshiba, & Aruga, 2007; Minocha et al., 2017; Rubin et al., 2010; Turrero Garcia et al., 2021; Wei et al., 2012). To determine the contribution of these progenitors to astrocytes in the septum, we generated a Zic4Cre; Sun1GFP reporter mouse line. Our staining indicated that LS, not MS astrocytes are mainly derived from Zic4-expressing progenitors (dLS: 77%; iLS:78%; vLS:71%; MS: 13%) **(Figure 1D and 1E)**. Similarly, Zic4-positive astrocytes in the LS make up less than 20% of total Zic4-derived cells (**Figure 1F**), consistent with previous observations, Zic4-Cre progenitors substantially contribute to other neuronal and glial cell types within the LS (Magno et al., 2022; Magno et al., 2017; Rubin et al., 2010; Turrero Garcia et al., 2021) (**Figure S1D-S1F**). To test the timing of astrogenesis within the septum and its associated progenitor niches, we stained for Sox9 with lineage markers in both wild-type (CD1) mice and astrocyte reporter line Aldh1l1Cre-EGFP at embryonic stages, and we observed that astrogenesis is already present in MS and LS respectively at as early as embryonic day (E)14, with MS astrocytes derived from Nkx2.1 progenitor enriched MGE and POA, while LS astrocytes originate from Zic progenitor enriched septum (**Figure S1G and S1H**), consistent with previous studies (Akdemir, Huang, & Deneen, 2020; Minocha et al., 2017; Molofsky et al., 2013). Taken together, our data suggests that MS and LS astrocytes are derived from distinct progenitor domains (**Figure 1G**). To investigate the morphology and tiling of MS and LS astrocytes, we used a GlastCreER; Ai14 reporter mouse line induced with tamoxifen at P10. Astrocytes in the MS can be distinguished from those in the LS by their longer branches (**Figures 1I-1K, S1I**), while the overall territory size and density of septal astrocytes did not significantly differ across subdivisions of the septum (**Figures 1H, 1L, and S1I**). To assess whether septal astrocyte tiling is refined during development, we examined the astrocyte morphology in MS and LS at P0, P3 and P8. Our data showed that astrocytes in both the MS and LS evenly distribute without distinct territorial boundaries segregating these regional astrocytes during early postnatal stages (**Figure S1J**). This suggests that astrocytes uniformly tile the septum regardless of the distinct anatomical regions they occupy. Further morphological analysis showed that MS astrocytes closely interact with neighbor parvalbumin-positive (PV+) neurons that are found in the MS, while LS astrocytes wrap adjacent Calbindin-positive (Calbindin+) neurons (**Figure S1K**), indicating distinct interactions between astrocytes and neurons in these anatomical regions. Neurons in the septum receive inputs from a diversity of excitatory and inhibitory projection neurons from other brain regions (Besnard & Leroy, 2022; Sheehan, Chambers, & Russell, 2004; Sweeney & Yang, 2015). To investigate interactions between astrocytes and specific types of synapses, we stained for the markers: vGlut1/PSD95 and vGAT/Gephyrin which identify glutamatergic and GABAergic synapses respectively, in P30 GlastCreER;Ai14 mice (**Figure 1M and 1N**). There were significantly fewer glutamatergic and GABAergic colocalized puncta within individual astrocyte territory in the MS compared to the LS (**Figure 1O and 1P**). This data collectively suggests that MS and LS astrocytes are derived from distinct developmental lineages and exhibit unique morphologies and synaptic interactions suited to the specific neuronal circuits they support.

**Figure 1:**
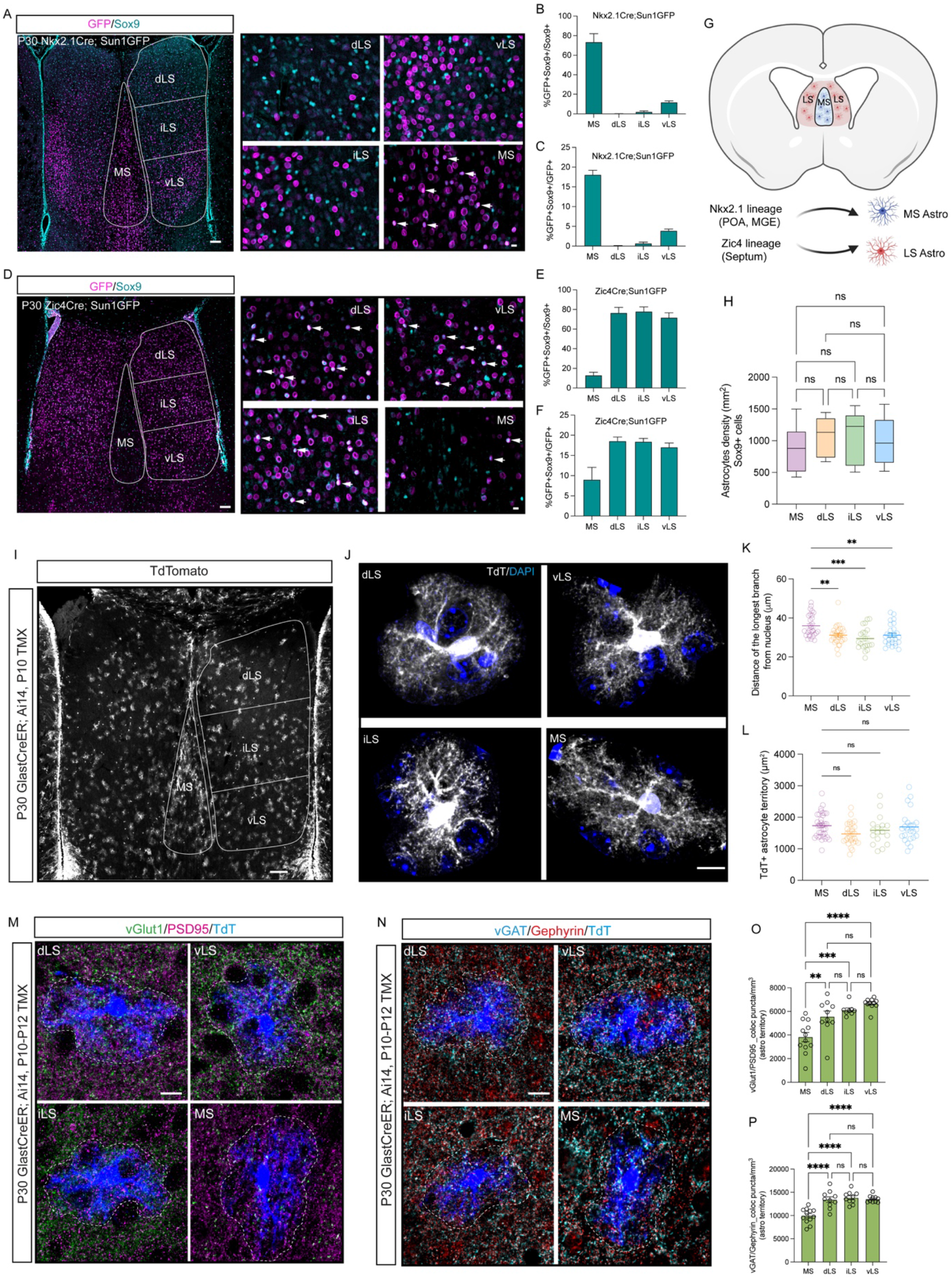
Astrocytes originated from Nkx2.1 and Zic4 lineages occupy MS and LS respectively. **A)** Immunostaining for GFP (magenta) and Sox9 (cyan) in P30 Nkx2.1Cre; Sun1GFP mice. Subdivisions of the septum are indicated with white lines. dLS: dorsal lateral septum; iLS: intermediate lateral septum; vLS: ventral lateral septum; MS: medial septum. White arrows indicate cells with overlapping signals. Scale bar: left: 100 µm, right: 10 µm. **B**) Proportion of Nkx2.1-derived astrocytes within the total astrocyte population. N=5 mice. **C)** Proportion of Nkx2.1-derived astrocytes in total Nkx2.1-derived cells. N=5 mice. **D**) Immunostaining for GFP (magenta) and Sox9 (cyan) in P30 Zic4Cre; Sun1GFP mice. Subdivisions of the septum are indicated with white lines. White arrows indicate cells with overlapping signals. Scale bar: left: 100 µm, right: 10 µm. **E**) Proportion of Zic4-derived astrocytes in total astrocytes. N=5 mice. **F)** Proportion of Zic4-derived astrocytes in total Zic4-derived cells. N=5 mice. **G**) Schematic of astrocyte lineages in the septum. Lateral ganglionic eminence (LGE); medial ganglionic eminence (MGE); preoptic area (POA). **H**) Astrocyte density in subdivisions of the septum. N=5 mice. **I**) Immunostaining for tdTomato (tdT+) (grey) in the septum of P21 GlastCreER; Ai14 mice injected intraperitoneally with tamoxifen at P10. Scale bar: 100 µm. **J**) High magnification of representative tdT+ astrocytes with DAPI staining in subdivisions of the septum. Scale bar: 10 µm. **K**) Analysis of the distance of the longest branch from the nucleus in different regions of the septum. N=4 mice, 20-30 astrocytes in each region. **L**) Analysis of the territories of tdT+ astrocytes (µm^2^). N= 4 mice, a total of 17-36 astrocytes in each region. **M**) Immunostaining for vGlut1 (green), PSD95 (magenta) and tdTomato (blue) in P21 GlastCreER; Ai14 septum, astrocyte territories are indicated with white dashed lines. Scale bar: 10 µm. **N**) Immunostaining for vGAT (cyan), Gephyrin (red) and tdTomato (blue) in P21 GlastCreER; Ai14 septum, astrocyte territories are indicated with white dashed lines. Scale bar: 10 µm. **O**) Quantification of excitatory synapse density in tdT+ astrocyte domains per mm^3^. N=3 mice, 8-12 astrocytes in each region. **P**) Quantification of inhibitory synapse density in tdT+ astrocyte domains per mm^3^. N=3 mice, 10-12 astrocytes in each region.

### Single-cell and spatial transcriptomics reveal astrocyte lineage-specific molecular profiles

The distinctive developmental origins and cellular features of MS and LS astrocytes imply that they may possess additional molecular specializations. To investigate this further, we collected the septa of Nkx2.1Cre; Sun1GFP mice at five developmental stages between P0 and P21 (**Figure 2A**). To distinguish cells with a history of Nkx2.1 expression from other lineages, we isolated GFP+ and GFP-nuclei at each time point and performed snRNA-Seq (**Figure 2A**). We collected a total of 166,418 nuclei (**Figure 2B)** from both Nkx2.1 (GFP+) and non-Nkx2.1 (GFP-) lineages (**Figure S2A**) across all time points (**Figure S2B**). A total of 22,221 astrocytic nuclei were classified into seven distinct subpopulations based on their transcriptional profiles (**Figures 2C, S2C and S2D**). These clusters were defined by both GFP expression (**Figure 2D and 2E**) and age (**Figures 2F, 2G and S2E**), where clusters 1, 5, and 6 primarily represented early astrocyte progenitors, clusters 0 and 3 represented LS astrocytes, and clusters 2 represented MS astrocytes. The clusters corresponding to astrocytes and neurons were the only ones exhibiting significant segregation based upon developmental origin (**Figure S2A and S2F**). To investigate the molecular differences between mature MS and LS astrocytes, we subclustered P21 astrocytes and identified three distinct clusters, including mature MS (GFP+) astrocytes (MSA), LS (GFP-) astrocytes (LSA), and one ventricular zone astrocyte progenitor cluster (LSP) (**Figure 2H, 2I and S2G**) characterized by high enrichment of Kif21a, Slit2 and Tmem132b (**Figure S2H**) (Cebrian-Silla et al., 2021; Desai, Velo, Yamada, Overman, & Engle, 2012; Y. Wang, Herzig, Molano, & Liu, 2022). The top 20 differentially expressed genes (DEGs) in mature MS and LS astrocyte populations revealed robust lineage-specific segregation in their molecular profiles, including the lineage-specific enrichment of Zic family members in GFP-astrocytes (**Figure 2H and 2J**). Higher-resolution clustering revealed additional subsets within the mature MS and LS astrocyte clusters (**Figure S2I**). Interestingly, these further segregated clusters did not exhibit significant differences in the top differentially expressed genes, suggesting that astrocyte molecular diversity is primarily defined by lineage (**Figure S2J-S2L**). Molecular similarities among astrocyte types are closely linked to their regional proximity to each other in the brain; thus, to further elucidate these relationships, we integrated our P21 snRNA-seq astrocytes with scRNA-seq data from cortical (CTX), hippocampal (HIP) and striatal (STR) astrocytes (Endo et al., 2022), which are proximal to septal astrocytes. Our data showed that both MS and LS astrocyte clusters are distinct from astrocytes in other regions based on their molecular signatures (**Figure S3A and S3B**). The majority of the top 10 differentially expressed genes in the septum were common to both MS and LS astrocytes, only Kcnd2 and Trpm3 were highly enriched in MS astrocytes (**Figure S3C**). Mapping shared genes from other brain regional astrocytes (Endo et al., 2022) to septal astrocytes revealed a high enrichment of those genes, although some genes were differentially expressed in either MS (e.g., *Slc4a4*, *Slc6a11*, *Slc39a12*, *Ndrg2*) or LS astrocytes (e.g., *Slc1a2* and *Prex2*) (**Figure S3D**). Further analysis showed that LS astrocytes expressed genes highly enriched in cortical and hippocampal astrocytes, while MS astrocytes exhibited an enrichment of genes specific to the midbrain/hindbrain (MB/HB) region (Endo et al., 2022) (**Figure S3E)**. This suggests a striking resemblance between septal astrocytes and those from other brain regions. To visualize the spatial distribution of astrocyte populations in the septum, we employed single-cell spatial transcriptomics (MERFISH) on the septum at P35, with a list of 500 genes enriched in defined astrocyte and neuron clusters. We identified 25 clusters including both astrocyte and neuron subpopulations, displaying differential spatial distributions in the P35 septum (**Figure S4A and S4B**). We selected astrocyte populations for further analysis using astrocyte markers *Aldh1l1* and *Slc1a2* (**Figure S4C**). We identified four astrocyte clusters, including LS, MS, and two ventricular zone astrocyte progenitor subgroups based on the expression of cluster-enriched genes (**Figures 2K-2M, S4D**), consistent with our snRNA-seq data (**Figures 2J and S2H**). Some genes exhibited expression gradients from high to low between LS and MS astrocytes (e.g. *Mfge8*, *Slc1a2*), and vice versa (e.g., *Lrig1*, *Slc6a11*) while others were exclusively expressed by LS (e.g., *Lhx2*, *Zic4*) or MS astrocytes at P35 (e.g., *Agt*, *Igst1*) (**Figure 2N and 2O**). Together, this data illustrates that MS and LS astrocytes exhibit distinct molecular profiles that correspond to their developmental origins.

**Figure 2:**
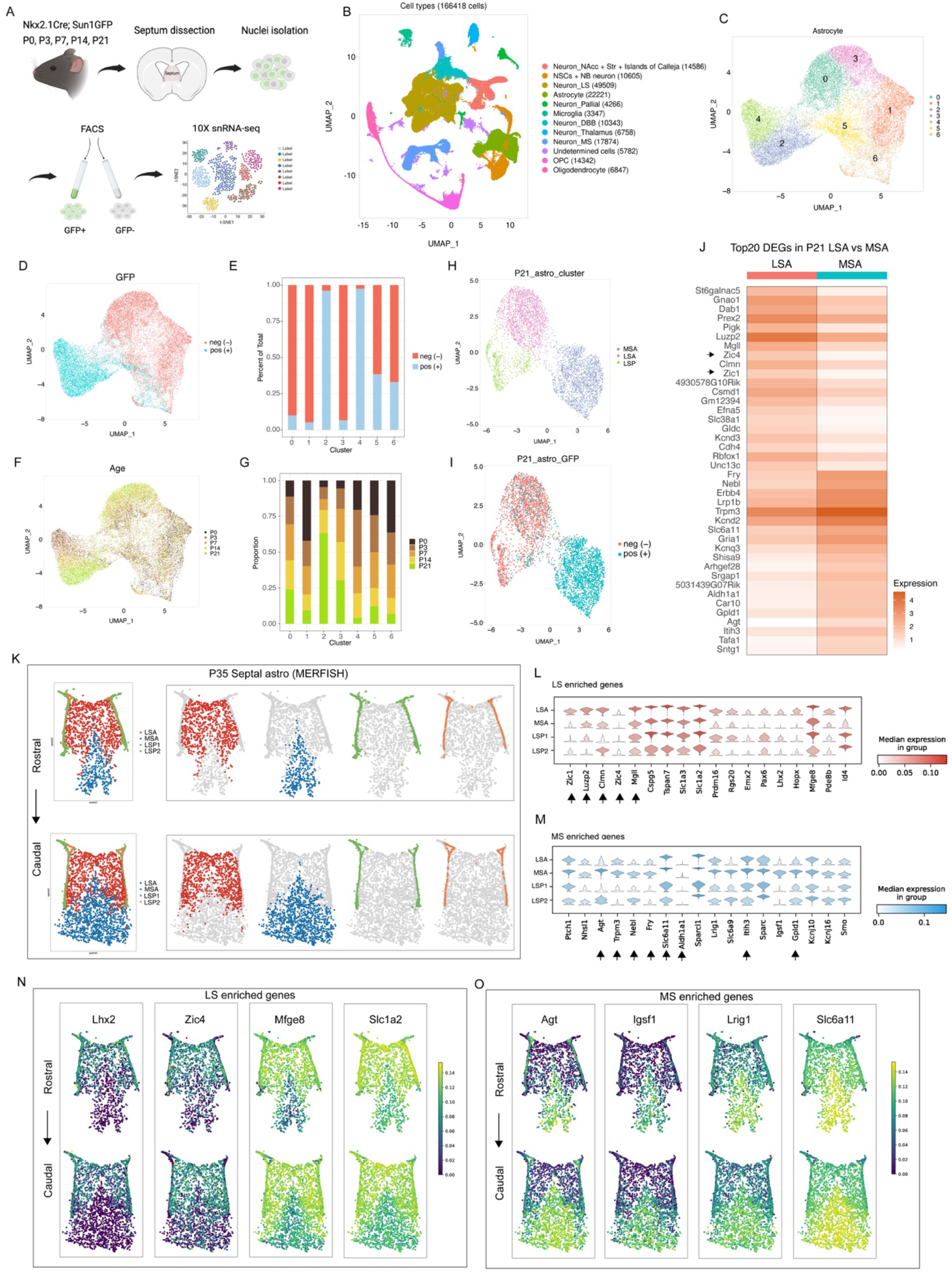
Lineage-specific astrocytes display distinct molecular signatures. **A)** Experimental design for snRNA-sequencing on sorted cells from Nkx2.1Cre; Sun1GFP septum at different developmental ages **B**) UMAP plot of cell types from all samples. **C**) UMAP plot of entire septal astrocyte clusters. **D**) UMAP plot of entire GFP+ and GFP-astrocytes. **E**) Proportion of GFP+ and GFP-astrocytes in each cluster. **F**) UMAP plot of septal astrocytes at individual ages. **G**) Proportion of cells in different clusters at each age. **H**) UMAP plot of P21 septal astrocytes at a 0.3 resolution level. MSA: medial septal astrocytes; LSA: lateral septal astrocytes; LSP: lateral septal astrocyte progenitors. **I**) UMAP plot of P21 GFP+ and GFP-astrocytes. **J**) Heatmap showing the top 20 differential expressed genes (DEGs) in LSA and MSA. Zic1 and Zic4 are highlighted. **K**) MERSCOPE analysis of P35 septal astrocyte clusters along the rostro-caudal axis. LSA: lateral septal astrocyte, MSA: medial septal astrocyte, LSP1: lateral septal astrocyte progenitor cluster 1, LSP2: lateral septal astrocyte progenitor cluster 2. **L, M**) MERSCOPE analysis showing gene enrichment in LS and MS astrocytes. The highlighted genes are present in the top 20 gene list in Figure 2J. **N, O**) MERSCOPE analysis showing representative genes enriched in MS and LS astrocytes along the rostro-caudal axis.

### Developmental trajectories of MS and LS astrocytes

Astrocyte development and maturation proceed through a series of transitional states influenced by intrinsic and extrinsic factors (Markey et al., 2023; Zarei-Kheirabadi et al., 2020). To investigate whether MS and LS astrocytes follow distinct molecular trajectories, we isolated MS and LS astrocyte populations across five developmental stages. Our analysis revealed clear and unique trajectories in both MS and LS astrocytes during their development (**Figure 3A and 3B**). To understand the transcriptional programs associated with the maturation of these regional astrocytes, we examined the top 10 temporally enriched genes for both MS and LS astrocytes (**Figure 3C and 3D**), and compared differential gene expression of MS versus LS astrocytes at P0-P14 (**Figures 3E-3H, S5A**). MS and LS astrocytes exhibited lineage-associated developmental gene expression programs. Genes such as *Sorcs3*, *Inka2*, *Cntnap5b*, *Car10* and *Lgi1* were highly enriched in developing MS astrocytes, while *Fstl5*, *Unc5d*, *Pigk*, *Hapln1* and *Mgll* were enriched in developing LS astrocytes (**Figure 3I and 3J**). To further assess the function of age-enriched genes, we performed gene ontology on septal astrocytes at each age. During early postnatal time points (P0-P3), gene ontology terms related to synapse organization and axonogenesis were significantly represented, while at P7, gene ontology terms were predominantly associated with metabolic processes (e.g., phospholipid, alcohol, and cholesterol synthesis). At later postnatal time points (P14-P21), gene terms were correlated with cell junction and synapse assembly (**Figure S5B)**. Differences in biological processes between MS and LS astrocytes were evident, with top genes enriched in P21 MS astrocytes linked to cell junction assembly and organization, and gene terms in LS astrocytes associated with dendrite development and glutamatergic synaptic transmission, consistent with glutamatergic inputs from the hippocampus into the septum (Khakpai, Zarrindast, Nasehi, Haeri-Rohani, & Eidi, 2013; Trent & Menard, 2010) (**Figure S5C**). Mapping gene families responsible for circuit-related functions (Endo et al., 2022) to our P21 astrocyte clusters, we identified differential expression patterns in the enrichment of K+ homeostasis and neurotransmitter transporters in both MS and LS astrocytes (**Figure S6A-S6E**). For instance, genes encoding K+ channels (*Kcnj10*, *Kcnj16*, *Kcnq3*, *Kcnd2* and *Atp1a2*) were highly expressed in MS astrocytes (**Figure S6A**). Neurotransmitter transporters, genes encoding glutamate transporters (*Slc1a2*, *Slc1a3* and *Slc1a4)* were highly enriched in LS astrocytes, while genes encoding GABA transporters (*Slc6a11* and *Slc6a1*) were highly enriched in MS astrocytes (**Figure S6E**), reflecting their interactions with different neural circuit inputs (Khakpai et al., 2013; Muller & Remy, 2018; Trent & Menard, 2010). Next, we used single-cell regulatory network inference and clustering (SCENIC) analysis (Aibar et al., 2017) to infer the transcription factor– target regulatory networks controlling lineage-specific and age-dependent astrocyte specification in the MS and LS. We identified 10 regulatory network clusters associated with both lineage and age (**Figure S7A-S8C**). Regulons such as *Zic4*, *Bach2*, *Prdm16*, *Pou2f1*, *Emx2* were enriched in LS astrocytes whereas *Nr2f1*, *Vezf1*, *Meis1*, *Smad3*, and *Dbx2* regulons were prominent in MS astrocytes (**Figure S7D-S7F**). SCENIC analysis revealed several genes enriched in LS astrocytes were inferred to be regulated by *Zic4*, including *Gnao1*, *Dab1* and *Efna5* (**Figure 2J**) (**Table S1**), while *Nr2f1* regulated genes like *Shisa9*, *Kcnd2*, *Slc6a11* were enriched in MS astrocytes. (**Table S1**). Moreover, *Tcf7l1* regulon was enriched at early time points (P0-P7), while *Bcl6* regulon was enriched at later time points (P14-P21) (**Figure S7G-S7I**). Certain regulons, including *Nr2f1*, *Vezf1* for MS and *Zic4*, and *Bach2* for LS, remain enriched throughout postnatal development (**Figure S7J**). These results suggest that septal astrocytes exhibit lineage-specific and age-dependent regulatory network enrichment. To further analyze the dynamic spatial and temporal expression patterns of region-specific genes in septal astrocytes, we studied the expression of *Slc6a11* and *Slc1a2* by *in situ* hybridization in the septum at P0, P3, P7, and P14. These genes respectively encode for astrocytic GABA and glutamate transporters that are essential for regulating neuronal circuit function (Ishibashi, Egawa, & Fukuda, 2019; Todd & Hardingham, 2020). We observed that differential expression of *Slc6a11* and *Slc1a2* increased between MS and LS astrocytes over time, with a comparable level in early stages that significantly diverged over development (**Figures 3K, 3L, 3N and 3O**), consistent with our snRNA-seq analysis, highlighting the refinement of gene expression in septal astrocytes by age (**Figure 3M and 3P**) and suggesting the potential influence of local environments on astrocyte molecular specification. Collectively, these data demonstrate that MS and LS astrocytes undergo unique lineage and region-specific differentiation that ultimately lead to their expression of specialized circuit-related functional gene programs.

**Figure 3:**
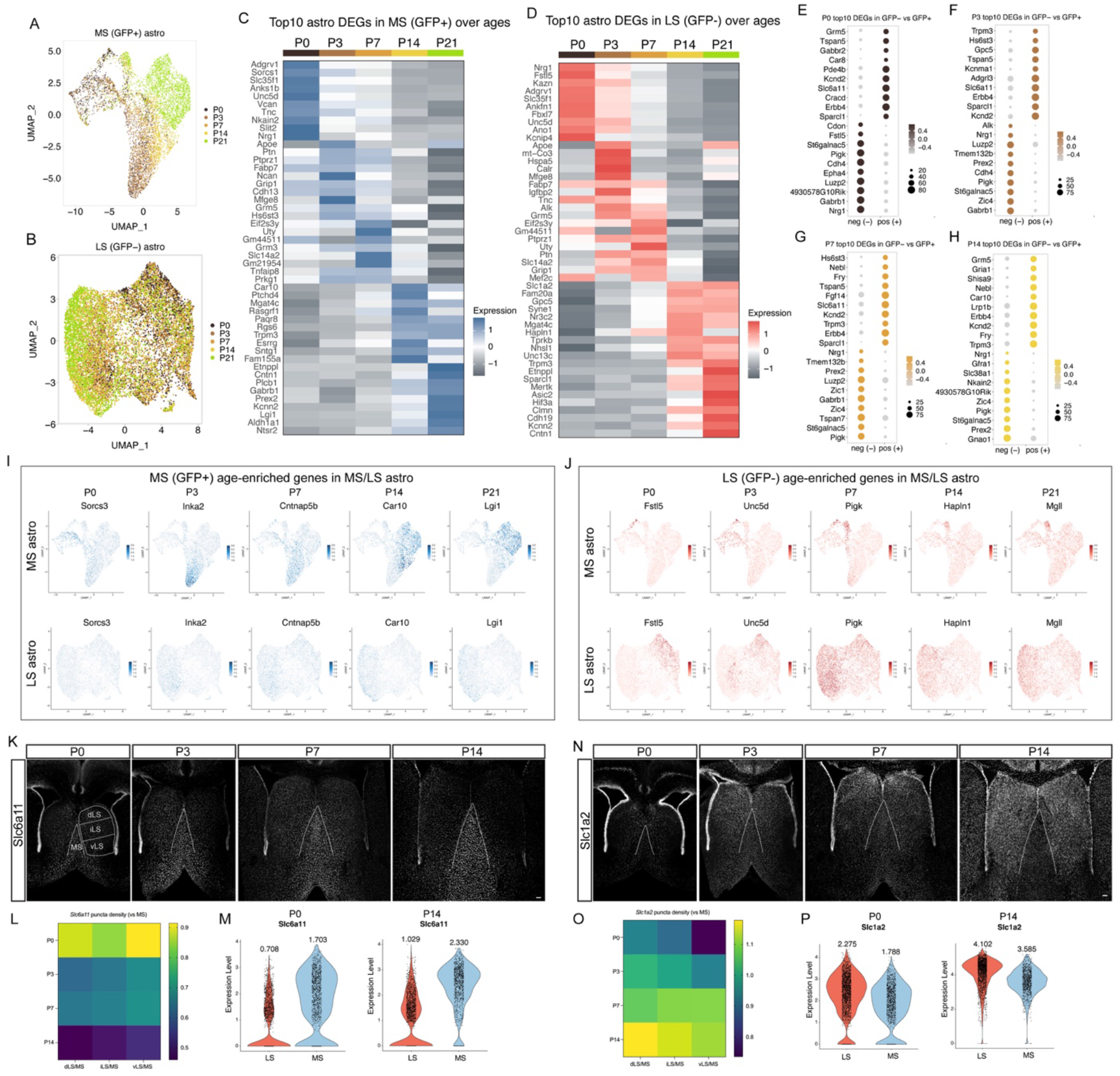
MS and LS astrocytes display unique age-dependent molecular profiles. **A, B)** UMAP plot showing age-dependent MS (**A**) and LS (**B**) astrocyte clusters. **C, D**) Heatmap showing the top 10 DEGs in GFP+ and GFP-astrocytes across ages. **E-H**) Heatmaps showing top 10 DEGs enriched in GFP+ vs GFP-at each age (P0-P14). **I**) Representative genes enriched in GFP+ astrocytes during development. **J**) Representative genes enriched in GFP-astrocytes during development. **K**) *Slc6a11* mRNA was detected in P0-P14 septum of wild-type (CD1) mice using *in situ* hybridization. Subdivisions of the septum are indicated with white lines. Scale bar: 100 µm. **L**) Heatmap showing the relative density of puncta of *Slc6a11* mRNA in each corresponding LS subnucleus, normalized by the density of puncta in the MS, at different ages. N=3-5 mice for each age. **M**) Violin plots showing *Slc6a11* expression (snRNA-seq) in both MS and LS astrocytes at P0 and P14. The average expression levels are indicated. **N**) *Slc1a2* mRNA was evaluated in P0-P14 septum of wild-type (CD1) mice using *in situ* hybridization. Scale bar: 100 µm. **O**) Heatmap showing the relative density of puncta of *Slc1a2* mRNA in each corresponding LS subnucleus, normalized by the density of puncta in the MS, at different ages. N=3-5 mice for each age. **P**) Violin plots showing *Slc1a2* expression (snRNA-seq) in both MS and LS astrocytes at P0 and P14. The average expression levels are indicated.

### Neuron-derived cues shape local astrocyte molecular identity

Astrocyte maturation is known to be influenced by crosstalk with neurons and other cell types in the local environment (Hill et al., 2019; L. M. Holt et al., 2019; Lanjakornsiripan et al., 2018; Scholze, Foo, Mulinyawe, & Barres, 2014; Xie et al., 2022). To ascertain which ligands and receptors participate in neuron-astrocyte interactions in each septal nucleus, we performed CellPhone DB (Efremova, Vento-Tormo, Teichmann, & Vento-Tormo, 2020) on defined neurons and astrocyte populations from both MS and LS (**Figures 4A, S8A-S8D**). Our data revealed that distinct ligand-receptor pairs were exclusively enriched in either MS or LS subsets of neurons and astrocytes (**Figures 4B, S8E-S8G**). For example, Bdnf-Ntrk2/Ncam1/Sort1 and Mdk-Ptprz1/Sorl1 pairs are highly expressed in subsets of MS neurons and astrocytes, while Eda-Edar and Fgf10-Fgfr1/2 pairs are exclusively enriched in subsets of LS neurons and astrocytes, suggesting unique interactions between specific subpopulations of neurons and astrocytes in the septum. Furthermore, we observed that several ligand-receptor pairs are not only region-specific but also age-dependent. For instance, the Mdk-Ptprz1 pair shows high enrichment at early ages, while the Bdnf-Ntrk2/Ncam1 pairs exhibit pronounced enrichment at later ages within the context of the MS (**Figure 4C**). Meanwhile, the Efnb2-Ephb2 pair show significant enrichment at early ages, while the Fgf10-Fgfr2 pair displays particularly robust enrichment at later ages within the LS (**Figure 4D**). Our previous research established that Shh, a ligand originating from deep-layer cortical neurons binds to Ptch1 expressed in cortical astrocytes. Shh signaling is necessary and sufficient to promote their layer-enriched molecular identity (Xie et al., 2022). Our snRNA-seq and MERFISH analyses show that Shh expression is highly enriched in subsets of MS neurons (**Figures 4E, 4G and S8E**), where expression increases over postnatal development (**Figure 4F**). Fgf family members exhibit enriched expression in LS neurons (**Figures 4E, 4H, S8F and S8H**), which increases with age (**Figure 4F**), implying that these neuron-derived ligands could contribute to region-specific astrocyte diversification. Notably, Shh receptor Ptch1 and Fgf receptor Fgfr3 are known to be highly enriched in astrocytes throughout the CNS (Pringle et al., 2003; Xie et al., 2022). We examined the expression of genes previously shown to be downstream of Shh and Fgf2 signaling in septal astrocytes (Lattke et al., 2021; Xie et al., 2022), and observed that a considerable proportion of these genes exhibit marked enrichment in the P21 MS and LS astrocytes, respectively (**Figure 4I and 4K**). *Lhx2*, a transcription factor downstream of Fgf2 signaling (Lattke et al., 2021), is exclusively expressed in LS astrocytes (**Figure S8I and S8J**). Downstream gene expression increased over time mirroring the temporal increase of Shh and Fgf ligand expression (**Figure 4J and 4L**). *Slc6a11*, a putative Shh pathway target gene (Xie et al., 2022) coding for a GABA neurotransmitter transporter that is essential for neural circuit function (Boddum et al., 2016; Minelli, DeBiasi, Brecha, Zuccarello, & Conti, 1996) is highly enriched in MS astrocytes (**Figure 2J and 2M**). To further validate whether *Slc6a11* expression is driven by Shh signaling pathway activation, we employed an astrocyte-specific conditional mutant mouse model wherein Shh signaling is hyperactivated by the conditional ablation of Ptch1 (Ptch1cKO) (Ferent et al., 2019; Xie et al., 2022). We observed that *Slc6a11* transcripts in LS were significantly increased in tdT+ astrocytes of Ptch1cKO mice compared to WT (**Figure 4M and 4N**). However, we did not observe a significant increase in MS astrocytes despite the loss of Ptch1, which could be attributed to saturating levels of Shh signaling already present in the MS. Collectively, our findings demonstrate that MS and LS astrocytes undergo regional subtype-specific signaling crosstalk that is exclusive to their local environments. Shh and Fgf function as two such neuron-derived cues that play important roles in shaping the specialized molecular identities of MS and LS astrocytes respectively.

**Figure 4:**
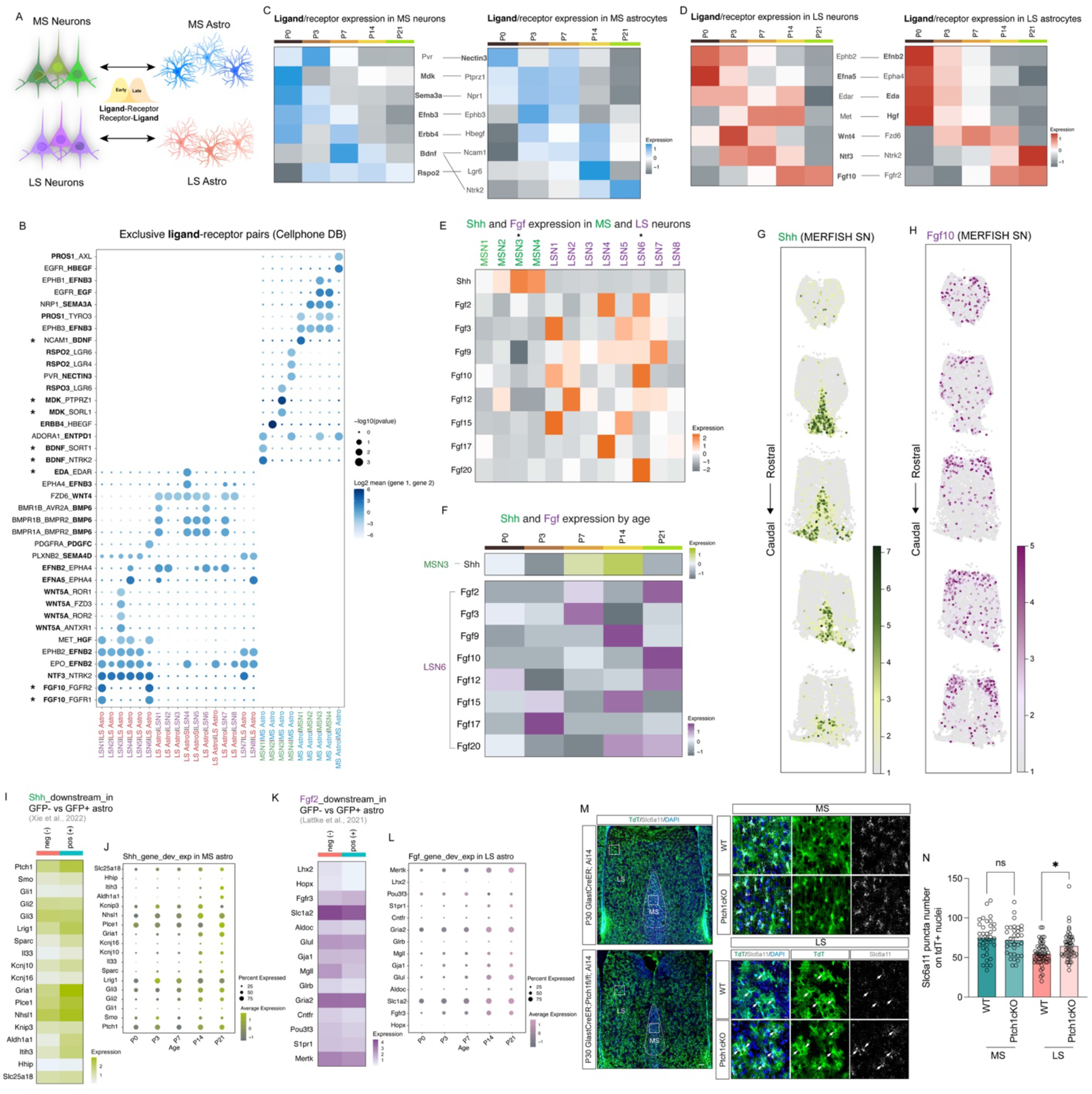
Region-specific and age-dependent interactions between subgroups of septal astrocytes and neurons. **A**) Diagram of putative astrocyte-neuron interactions during development. **B**) CellphoneDB analysis reveals exclusive ligand-receptor pairs between subclusters of lateral septum neurons (LSN) (purple) with LS astrocytes (red), and subclusters of medial septum neurons (MSN)(green) with MS astrocytes (blue). The highlighted pairs indicate representative ones present in subsets of neuron and astrocyte interactions. **C**) Age-dependent ligand-receptor pairs in MS neurons and astrocytes. Ligands are highlighted in bold. **D**) Age-dependent ligand-receptor pairs in LS neurons and astrocytes. Ligands are highlighted in bold. **E**) Heatmap showing that Shh and Fgfs are highly enriched in MSN and LSN subclusters. The highlighted subclusters indicate the ones with the most enriched Shh and Fgfs. **F**) Heatmap showing an increase in the expression of Shh and Fgfs in MSN3 and LSN6 subclusters respectively. **G**) MERSCOPE showing Shh expression patterns in P35 septal neurons through the rostral and caudal axis. SN: septal neuron. **H**) MERSCOPE showing Fgf10 expression patterns in P35 septal neurons through the rostral and caudal axis. SN: septal neuron. **I**) Heatmap showing selected Shh downstream genes are specifically enriched in P21 MS astrocytes. **J**) Dotplot showing increases in Shh downstream gene expressions during development. **K**) Heatmap showing selected Fgf2 downstream genes are specifically enriched in P21 LS astrocytes. **L**) Dotplot showing increases in Fgf2 downstream gene expressions during development. **M**) *In situ* hybridization of *Slc6a11* combined with immunostaining for tdTomato in P21 tamoxifen-induced GlastCreER;Ai14 (WT) and GlastCreER; Ptch1fl/fl; Ai14 (Ptch1cKO). White boxes indicate the selected regions for high magnification images (right panels). White arrows indicate the recombined astrocytes. Scale bar: left: 100 µm, right: 10 µm. **N**) Quantification of *Slc6a11* puncta number on tdT+ astrocyte nuclei. N=4 mice, 33-59 astrocytes in each condition.

### Transplanted astrocytes adopt local astrocytic morphology and molecular identity

Astrocyte molecular identities are thought to arise through a combination of developmentally determined factors related to their lineage origins and extrinsic cues from their local environment (Markey et al., 2023; Zarei-Kheirabadi et al., 2020). However, the relative contribution of intrinsic programming versus environmental factors is not fully determined. To address these questions, we performed homotopic and heterotopic transplantations of fluorescently labeled Nkx2.1-derived MS and Zic4-derived LS cells of P3-P5 mice into the septal region of wild-type mice of the same age (**Figure 5A**). We examined the morphology and gene expression of transplanted cells at P30, a time point at which the majority of cells had differentiated into astrocytes and oligodendrocytes, as neuronal survival throughout the procedure was limited (**Figure 5B**). Heterotopically transplanted Nkx2.1-derived astrocytes within the LS exhibited a shorter branch pattern compared to homotopically transplanted cells, and more closely resembled local endogenous LS astrocytes (**Figures 5C, 5D and 1K**). Similarly, grafted Zic4-derived astrocytes in the MS displayed elongated branches compared to homotopically transplanted cells (**Figures 5E, 5F, and 1K**). To ascertain whether transplanted astrocytes adopted local astrocyte molecular identities, we probed the expression of *Slc1a2* and *Slc6a11* transcripts, normally enriched in LS and MS respectively (**Figure 2N and 2O**) in the graft-derived astrocytes. Nkx2.1-derived astrocytes within the LS exhibited an upregulation of *Slc1a2* transcripts and a downregulation of *Slc6a11* transcripts (**Figure 5G-5J**). Conversely, *Slc1a2* transcripts were decreased while *Slc6a11* transcripts were increased in Zic4-derived astrocytes transplanted into the MS when compared to homotopic transplants (**Figure 5K-5N**). Additionally, we found that Lhx2 levels were increased in Nkx2.1-derived astrocytes within the LS and diminished in Zic4-derived astrocytes within the MS (**Figure S9A-S9D**). However, these expression patterns did not perfectly mirror those of the endogenous local astrocytes, as homotopically transplanted LS astrocytes had reduced Lhx2 compared to resident astrocytes, suggesting this reduction could be due to the transplantation procedure itself. Taken together, our findings strongly suggest that extrinsic cues play a pivotal role in shaping the ultimate molecular and morphological identities of septal astrocytes, such that cells from a different lineage origin can still adopt functional features consistent with the local environment.

**Figure 5:**
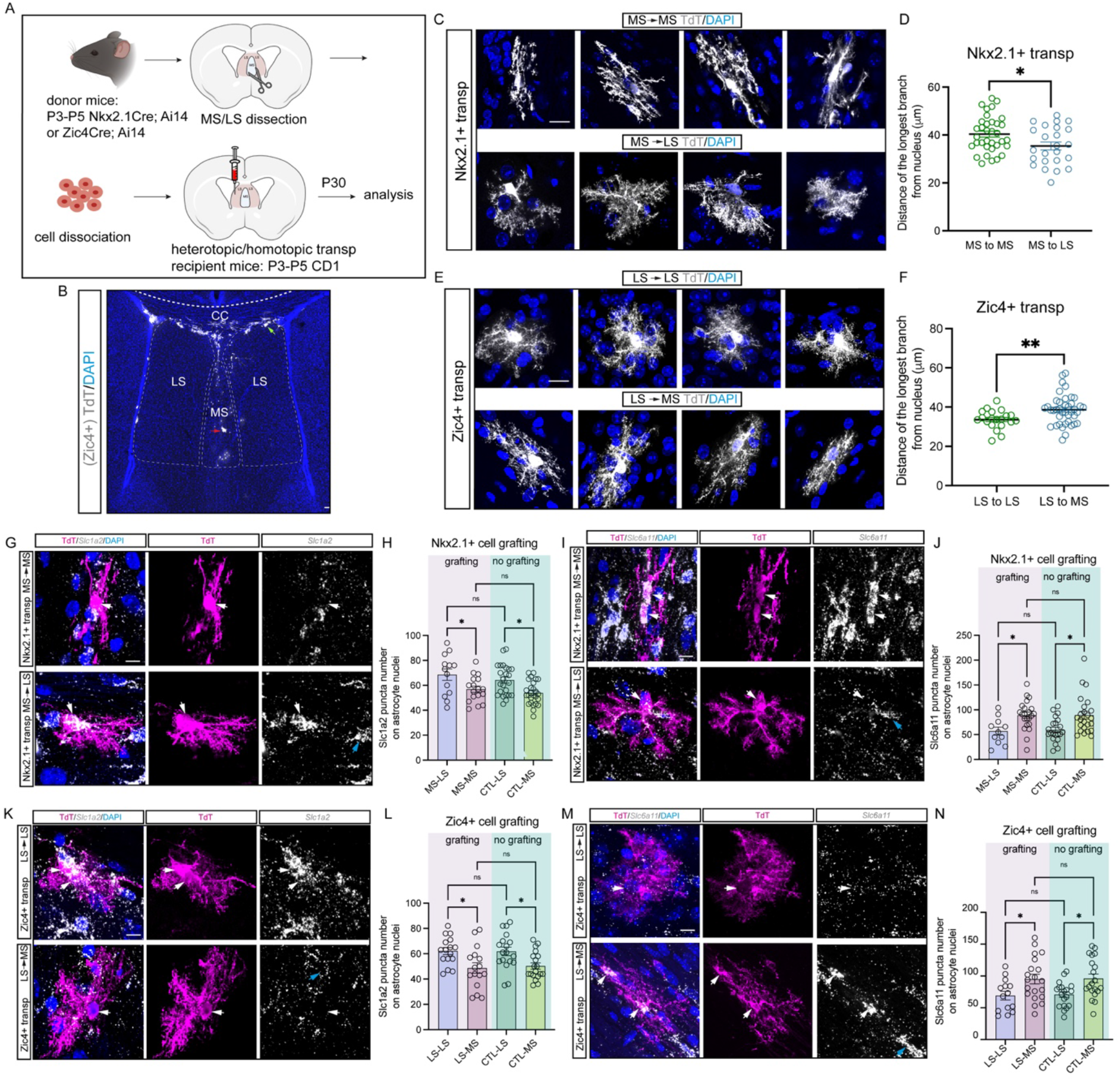
Grafted astrocytes acquire local astrocyte morphology and molecular identities. **A**) Experimental design for cell transplantation experiments in the septum. **B**) Representative image of P30 Zic4-derived cells when grafted into the MS at P4. Immunostaining for tdTomato (grey) and DAPI (blue). The red and green arrows indicate grafted cells in MS and LS respectively. Scale bar: 100 µm. **C**) Immunostaining for tdTomato and DAPI in grafted Nkx2.1-derived astrocytes in heterotopic (MS to LS) and homotopic (MS to MS) conditions. Scale bar: 10 µm. **D**) Quantifications of the length of the longest branch in grafted astrocytes in MS to LS compared to MS to MS groups. N= 5-9 transplants, 24-35 astrocytes were analyzed for each condition. **E**) Immunostaining for tdTomato and DAPI in grafted Zic4-derived astrocytes in heterotopic (LS to MS) and homotopic (LS to LS) conditions. Scale bar: 10 µm. **F**) Quantifications of the length of the longest branch in grafted astrocytes in LS to MS groups compared to LS to LS groups. N= 4-6 transplants, 20-40 astrocytes were analyzed for each condition. **G**) *In situ* hybridization of *Slc1a2* combined with tdTomato staining in transplanted Nkx2.1-derived astrocytes. White arrows indicate transplanted astrocytes; blue arrow indicates a host astrocyte. Scale bar: 10 µm. **H**) Quantification of *Slc1a2* transcripts on astrocyte nuclei in both grafting and no grafting groups. N=4-6 transplants, 13-26 astrocytes were analyzed for each group. **I**) *In situ* hybridization of *Slc6a11* combined with tdTomato staining in transplanted Nkx2.1-derived astrocytes. White arrows indicate transplanted astrocytes; blue arrow indicates a host astrocyte. Scale bar: 10 µm. **J**) Quantification of *Slc6a11* transcripts on astrocytes nuclei in both grafting and no grafting groups. N=4-6 transplants, 11-24 astrocytes were analyzed for each group. **K**) *In situ* hybridization of *Slc1a2* combined with tdTomato staining in transplanted Zic4-derived astrocytes. White arrows indicate transplanted astrocytes; blue arrow indicates a host astrocyte. Scale bar: 10 µm**. L**) Quantification of *Slc1a2* transcripts on astrocytes nuclei in both grafting and no grafting groups. N=4-6 transplants,16-19 astrocytes were analyzed for each group. **M**) *In situ* hybridization of *Slc1a2* combined with tdTomato staining in transplanted Zic4-derived astrocytes. White arrows indicate transplanted astrocytes; blue arrow indicates a host astrocyte. Scale bar: 10 µm**. N**) Quantification of *Slc1a2* transcripts on astrocyte nuclei in both grafting and no grafting groups. N=4-6 transplants, 14-20 astrocytes were analyzed for each group.

## DISCUSSION

In this study, we show that astrocytes allocated to the MS and LS have distinctive progenitor origins, molecular profiles and morphologies suited to their specialized roles within each subregion. Our genetic fate mapping revealed that MS astrocytes are derived from Nkx2.1-expressing progenitors and most likely derive from Nkx2.1 expressing region of the MGE and POA (Minocha et al., 2017; Turrero Garcia et al., 2021; Wei et al., 2012), while LS astrocytes are produced within the septal proliferative zone by progenitors that express Zic family transcription factors, including Zic4 (**Figure S1H**). However, whether intrinsic or extrinsic mechanisms underlie astrocyte specification is not understood. The sharp segregation of Nkx2.1 versus Zic4-derived astrocytes might suggest that many of the distinctive features of MS and LS astrocytes are genetically encoded through their lineage. However, our data supports a model where the local cellular environment substantially contributes to the final specification of astrocytes. Our transplantation experiments show that extrinsic factors are sufficient to induce astrocytes to adopt both the morphological and molecular features of their local environment. However, intrinsically encoded factors related to lineage origins likely play important roles in the specification of septal astrocytes, such as guiding the migration and positioning of astrocyte precursors into the lateral or medial septum. However, once cells reach their final positions, local factors appear to take over, as the complement and concentration of extrinsic signaling cues dictate the final specification of astrocytes. This type of mechanism fits with the overall view of astrocytes as sensors of local environmental conditions with the capacity to adjust to them in order to maintain homeostasis and support circuit adaptations. This model aligns with the cellular and molecular plasticity we observed in our heterotopic transplantations (**Figure 5**). In the future, it will be important to determine the constraints on astrocyte plasticity that may be dictated by lineage origins or other factors, and whether there is a critical developmental window during which astrocytes can adopt region-specific features. This will have important implications for the utilization of astrocytes in cell-based therapies.

The complex morphologies of astrocytes are essential for the performance of their diverse functions in the brain allowing them to interact with a range of cell types, including neurons, other glia and blood vessels. MS astrocytes exhibit elongated branches that align to axon fibers, similar to fibrous astrocytes found in white matter regions (Akdemir et al., 2020; Rowitch & Kriegstein, 2010). This alignment suggests a potential influence of the dense fibers that traverse the MS in shaping the morphology of adjacent astrocytes. However, the specific factors originating from the local circuitry that contribute to this distinctive astrocyte morphology remain largely unidentified. Recent research efforts have explored the gene networks related to astrocyte morphology across a diverse range of brain regions and disease models (Endo et al., Science, 2022). Their findings suggest a complex interplay between the anatomical structure of local neurons and astrocyte-autonomous molecules in shaping astrocyte morphology. Future research should address the precise molecular interactions governing astrocyte morphogenesis within the septum, providing insight into both the diverse forms of astrocytes within this specific region and the broader spectrum of astrocyte morphologies present in diseases associated with astrocytic morphological defects.

Our snRNA-seq data reveals that MS and LS astrocytes exhibit lineage-specific molecular identities during development and highlights two critical periods of astrocyte molecular specification: P0-P3, a phase marked by axonogenesis and the initiation of neuron wiring, and P7-P14, a peak period of synapse formation, assembly, and pruning (Farhy-Tselnicker & Allen, 2018) (**Figures 3 and S5**). Eph receptors and their ephrin ligands, well-known for their functions in contact-dependent communication between cells and axon guidance (Egea & Klein, 2007; Feldheim & O’Leary, 2010; Murai & Pasquale, 2011; Pasquale, 2005; Wilkinson, 2001) are highly enriched in both MS and LS neurons and astrocytes, with Ephb3 is expressed in MS astrocytes, and its receptor Efnb3 expressed in MS neurons, while Epha4 and Efnb2 are expressed in LS astrocytes and their corresponding ligand and receptor, Efna5 and Ephb2, are highly enriched in LS neurons, particularly at early developmental stages. This phenomenon aligns with the role of specialized GFAP+ midline glial populations in cortical development which express guidance molecules influencing axon targeting (Shu, Puche, & Richards, 2003; Shu & Richards, 2001; Unni et al., 2012). In later postnatal stages, Brain-derived neurotrophic factor (BDNF) and neurotrophin 3 (NTF3) are exclusively and robustly enriched in MS and LS neurons respectively, while their receptors Ntrk2 (TrkB) are enriched in both MS and LS astrocytes. BDNF-Ntrk2 (TrkB) pathway has been shown to be important for mediating and regulating astrocyte morphology, synaptogenesis, and neuronal activity (Cunha, Brambilla, & Thomas, 2010; Fernandez-Garcia et al., 2020; L. M. Holt et al., 2019; Rodriguez et al., 2023). This illustrates that while MS and LS astrocytes exhibit many shared molecular features, they could acquire the capacity to differentially respond to particular ligands from neighboring neurons. Our data shows that astrocyte-neuron interactions are not only age-dependent, but also region and subgroup-specific. For example, interactions such as Fgf10-Fgfr2, Eda-Edar, Hgf-Met are exclusively expressed in LS astrocyte-neuron interactions (**Figure 4B**), with corresponding ligands and receptors enriched in subsets of LS neurons (**Figure S8F**). Similarly, pairs like Bdnf-Ntrk2 (TrkB) and Rspo2-Lgr6 are selectively expressed in MS interactions (**Figure 4B**), with ligands enriched in subsets of MS neurons (**Figure S8E**). These observations suggest the presence of unique interactions within specific subsets of neurons and astrocytes in the septum. Shh, a morphogen derived from neurons, signals to astrocytes expressing its receptor Ptch1 thereby regulating their molecular and functional profiles in the cerebellum and cortex (Farmer et al., 2016; Garcia, Petrova, Eng, & Joyner, 2010; Xie et al., 2022). Here, the Shh signaling pathway strongly impacts septal astrocyte molecular identity highlighting the functional conservation of this ligand-receptor pair across different brain regions. Moreover, our data unveiled several other critical ligand-receptor pairs that are likely to play a role in astrocyte specification, underscoring the extensive array of molecules involved in neuron-astrocyte interactions. These findings raise intriguing questions about the cooperative functional roles of these ligand-receptor pairs, especially in the context of age, cellular types, and specific brain regions where these interactions take place. Exploring these aspects in future studies could provide essential insights into the precise molecular codes governing intercellular communication between astrocytes and other cells in the brain.

Medial and lateral nuclei of the septum are distinguished by their patterns of connectivity with the hippocampus, where neurons in the MS project extensively to the hippocampus and receive inputs from a subset of GABAergic neurons in the same region (Muller & Remy, 2018; Sun et al., 2014). On the other hand, LS neurons do not project directly to the hippocampus but receive substantial inputs from glutamatergic neurons located in the CA region (Besnard & Leroy, 2022; Risold & Swanson, 1997; Sheehan et al., 2004). These differences in the circuitry are reflected in the molecular profiles of MS and LS astrocytes, where MS astrocytes are highly enriched for GABA transporter genes (*Slc6a11*, *Slc6a1*), while LS astrocytes show enrichment for glutamate transporter genes (*Slc1a2*, *Slc1a3*, *Slc1a4*) (**Figure S6E**). There is evidence that glutamate and GABA release induce transcriptional, cellular and functional changes in astrocytes (Cheng et al., 2023; Duan, Anderson, Stein, & Swanson, 1999; Morel, Higashimori, Tolman, & Yang, 2014). However, whether glutamatergic and GABAergic neurotransmitter release is entirely responsible for transporter specialization is not entirely clear. Our data suggests that neuron-derived factors such as Shh and Fgf play important roles in astrocyte functional specialization in the septum. We show that astrocyte-specific activation of Shh signaling is sufficient to promote the upregulation of *Slc6a11* in LS astrocytes (**Figure 4M and 4N**). We also observe the increased expression of *Slc1a2* and *Lhx2*, downstream targets of Fgf signaling in heterotopic transplanted LS astrocytes (Lattke et al., 2021) (**Figures 5G, 5H, S9A and S9B)**. These findings support our view that neuron-derived cues are essential for the expression of functional genes that specialize astrocytes for their circuit-specific roles.

Collectively, our data defines the molecular heterogeneity of astrocytes within the septum and uncovers the mechanisms that underlie astrocyte specifications during septal development. Recent technological advancements have revealed novel functions of astrocytes in both health and disease, shedding light on their potential as therapeutic targets. Understanding the intricate interactions between astrocytes and other cell types during development may provide insights into astrocyte regulation during circuit formation and refinement, thereby inspiring new ideas for therapeutic interventions (Clark et al., 2021; Lee, Wheeler, & Quintana, 2022; Stogsdill, Harwell, & Goldman, 2023).

## AUTHOR CONTRIBUTIONS

Conceptualization, Y.X. and C.C.H.; Investigation, Y.X., C.M.R., A.A.G., M.T.G., F.D.H., S.M.H., W.M., J.L., M.A., O.M.; Resources, A.O.P., A.A.B. and C.C.H.; Writing, Y.X. and C.C.H.; Editing, Y.X., C.M.R., A.A.G., M.T.G., C.C.H.; Funding Acquisition, C.C.H.; Supervision, C.C.H.

## ACKNOWLEDGEMENTS

The authors would like to thank all members of the Harwell lab for feedback and support; Rhiana Simon, Yuqi Ren and Denise Garcia for comments on the manuscript; This research was supported by NIH Grants R01MH119156 and R01NS11893 to CCH.

## KEY RESOURCES TABLE

| REAGENT or RESOURCE | SOURCE | IDENTIFIER |
| --- | --- | --- |
| **Antibodies** |  |  |
| Rabbit monoclonal anti-Lhx2 | Millipore | ABE1402 |
| Rabbit monoclonal anti-RFP | Rockland | 600-401-379 |
| Goat monoclonal anti-Sox9 | R&D | AF3075 |
| Rabbit monoclonal anti-Olig2 | Millipore | AB9610 |
| Rabbit polyclonal anti-Zic | A gift from Segal Lab, DFCI | Borghesani et al., 2002 |
| Mouse monoclonal anti-Nkx2.1 | Santa Cruz | 53136 |
| Chicken polyclonal anti-GFP | Aves | Cat # GFP-1020 |
| Chicken polyclonal anti-RFP | Rockland | Cat # 600-901-379 |
| Rabbit monoclonal anti-PSD95 | Thermo Fisher | 51-6900 |
| Guinea pig monoclonal anti-VGluT1 | Millipore | AB5905 |
| Guinea pig monoclonal anti-VGAT | synaptic systems | 131308 |
| Rabbit monoclonal anti-Gephyrin | synaptic systems | 147008 |
| Goat polyclonal anti-chicken Alexa 488 | Thermo Fisher | Cat # A11039 |
| Goat polyclonal anti-chicken Alexa 546 | Thermo Fisher | Cat # A11040 |
| Goat polyclonal anti-Guinea pig Alexa 488 | Thermo Fisher | Cat # A11073 |
| Goat polyclonal anti-mouse Alexa 488 | Thermo Fisher | Cat # A11001 |
| Goat polyclonal anti-mouse Alexa 647 | Thermo Fisher | Cat # A21236 |
| Goat polyclonal anti-rabbit Alexa 488 | Thermo Fisher | Cat # A11034 |
| Goat polyclonal anti-rabbit Alexa 647 | Thermo Fisher | Cat # A21245 |
| Donkey polyclonal anti-goat Alexa 647 | Thermo Fisher | Cat # A21447 |
| Donkey polyclonal anti-mouse Alexa 488 | Thermo Fisher | Cat # A21202 |
| **RNAscope Probes** |  |  |
| Slc6a11 | ACD | 492661-C3 |
| Slc1a2 | ACD | 441341 |
| **Critical Commercial Assays** |  |  |
| MERSCOPE | Vizgen | 91600001 Rev C |
| Chromium next GEM single cell 3'reagent kits v3.1(dual index) | 10x Genomics | CG000315 Rev C |
| RNAscope Fluorescent Multiplex Assay Part 1 and Part 2 | ACD | Document # 320513-USM;<br>Document # 320293 |
| **Experimental Models:**<br>**Organisms/Strains** |  |  |
| Mouse: CD1 | Charles river laboratories | Strain code 022 |
| Mouse: C57BL/6J-Tg(Nkx2-1-cre)2Sand/J (Nkx2.1-Cre) | The Jackson Laboratory | Stock No: 008661 |
| Mouse: B6;129-Gt(ROSA)26Sor<sup>tm5(CAG-Sun1/sfGFP)</sup>Nat/J | The Jackson Laboratory | Stock No: 021039 |
| Mouse: Zic4Cre | Kessaris lab, UCL | Rubin et al., 2010 |
| Mouse: B6.Cg-<br>*Gt(ROSA)26Sor<sup>tm14(CAG-tdTomato)Hze</sup>/J* (Ai14) | The Jackson Laboratory | Stock No: 007914 |
| Mouse: Tg(Slc1a3-cre/ERT)1Nat/J<br>(Glast-CreER<sup>T2</sup>) | The Jackson Laboratory | Stock No: 012586 |
| Mouse: B6N.129-*Ptch1<sup>tm1Hahn</sup>/J*<br>(*Ptch1<sup>flox/flox</sup>*) | The Jackson Laboratory | Stock No: 012457 |
| Mouse: Tg(Aldh1l1-EGFP,-<br>DTA)D8Rth/J | The Jackson Laboratory | Stock No: 026033 |
| **Software and Algorithms** |  |  |
| Fiji 2.0.0-rc-69/1.52p | (Schindelin et al., 2012) | https://imagej.net/Fiji |
| Prism 9 | GraphPad |  |
| MATLAB R2019b | Mathworks | https://www.mathworks.com/products.html |
| R studio |  | http://www.rstudio.com/ |
| Spyder |  | https://www.spyder-ide.org/ |

## LEAD CONTACT AND MATERIAL AVAILABILITY

This study did not generate new unique reagents. Further information and requests for resources and reagents should be directed to and will be fulfilled by the Lead Contact, Corey Harwell.

## EXPERIMENTAL MODEL AND SUBJECT DETAILS

All animal procedures conducted in this study followed experimental protocols approved by the Institutional Animal Care and Use Committee of Harvard Medical School (HMS) and the University of California, San Francisco (UCSF). Mouse lines are listed in the key resources table. Mouse housing and husbandry conditions were performed in accordance with the standards of the Center of Comparative Medicine at HMS and the Laboratory Animal Resource Center (LARC) at UCSF. Embryonic (E) day 14, E17 and Postnatal (P) day 1-35 mice were used for this study. The embryonic and postnatal mouse ages for each experiment are indicated in figures and figure legends.

## METHOD DETAILS

### Tamoxifen administration

Tamoxifen (Sigma) was dissolved in corn oil at a concentration of 20 mg/ml at 37°C and stored at 4°C for the duration of injections. For mice injected at P10 (WT: Glast-CreERT2; Ai14 or Ptch1cKO: Glast-CreERT2; Ptch1flox/flox; Ai14), 10-15 µL of tamoxifen was injected intraperitoneally for three consecutive days in order to induce efficient recombination. For astrocyte morphology analysis, we injected a single dose into Glast-CreERT2;Ai14 mice to sparsely label astrocytes.

### Immunohistochemistry (IHC)

Postnatal animals were transcardially perfused with PBS followed by 4% paraformaldehyde (PFA), their brains were dissected out and post-fixed in 4% PFA overnight at 4°C. Brains were sliced into 50 µm sections on a vibratome (Leica Microsystems VT1200S). Sections were prepared and placed in blocking solution (0.3% Triton (Amresco), and 10% goat serum in PBS) for 1-2 h at room temperature. Followed by primary antibodies overnight at 4°C and secondary antibodies for 1 h at room temperature (RT). DAPI (4’,6-diamidino-phenylindole, Invitrogen) was added to the secondary antibody solution. The sections were mounted using ProLong Gold Antifade Mountant (Invitrogen). Cryosections: Brains were cryoprotected in 30% sucrose/PBS overnight at 4°C after post-fixation. Brains were embedded in O.C.T. compound (Sakura), frozen on dry ice and stored at -80°C. Samples were sectioned at 20 µm on a cryostat (ThermoFisher CryoStart NX70). Images were acquired using a Leica SP8 or STELLARIS laser point scanning confocal microscope,10X, 20X, 40X, 63X and 100X objectives were used and images were further analyzed using Fiji. Brightness and contrast were adjusted as necessary for visualization, but the source images were kept unmodified. For Astrocyte morphology analyses, Z-stack images were collected to maximize the coverage of the entire astrocyte territory. The territory and the length from the nucleus to the end of the longest branch were measured in Fiji.

### Single-nucleus RNA-sequencing (snRNA-seq) and analysis

The septum was dissected out from Nkx2.1Cre; Sun1GFP mice. Nuclei were isolated from septal tissue. The number of biological replicates used was as follows: for P0 and P3, two pooled samples were used (each pool contained 3 mice; their sex was not determined); for P7, two males were used; for P14: one female and one male; and for P21: two females and two males were used. Both GFP+ and GFP-nuclei were sorted and counted for cell density, followed by 10X Genomics platform (Chromium next GEM single cell 3’reagent kits v3.1(dual index)) and sequencing using the NovaSeq 6000 sequencer. Sequence reads were processed and aligned to the mouse geneome GRC38 USING 10X Genomics Cell Ranger 6.0.0. Nuclei with fewer than 200 genes, total counts less than 1000, and a percentage of mitochondrial reads greater than 5% were excluded from the analysis (for astrocyte cluster analysis, mitochondrial reads greater than 2% of total reads were excluded from the analysis). For snRNA-seq analysis at different ages, a total of 166,418 nuclei (P0: 35,066 nuclei; P3: 36,692 nuclei; P7: 29,034 nuclei; P14: 19,926 nuclei; P21: 45,700 nuclei; GFP+: 72,573 nuclei; GFP-: 93,845 nuclei) were used for downstream analysis. Low read counts were normalized, and dimensional reduction was performed by PCA (n = 50), followed by regressing out percent mitochondrial reads and number of genes per nucleus. Data from different developmental stages was integrated using the Seurat 4 anchor-based integration pipeline with reciprocal PCA, as recommended by the authors when integrating samples with potentially non-overlapping populations. Cell types are determined by examining the expression of canonical marker genes. For differential expression analysis, the Seurat object was generated and followed by the FindMarkers() function. Gene ontology enrichment analysis was performed using the enrichGO() function. Cortical, hippocampal, and striatal scRNA-seq data were obtained from a previously published dataset (Endo et al., 2022) and integrated with P21 snRNA-seq septal astrocytes and normalized for following analysis. Fgf2 downstream targets are shown previously (Lattke et al., 2021). Genes that were both Fgf2 downstream targets and highly enriched in LS astrocytes were selected in Figure 4K. Shh downstream targets were shown previously (Xie et al., 2022). Genes that were both Shh downstream targets (cut off p<0.01) and highly enriched in MS astrocytes, plus known Shh signaling pathway members (Ptch1, Smo, Gli family members) (Carballo, Honorato, de Lopes, & Spohr, 2018) were selected in Figure 4I. Cell-cell interaction analysis between different cell populations was performed using CellphoneDB version 3.0 and method=’statistical_analysis’ (Efremova et al., 2020). The ligand-receptor pairs expressed in both MS and LS neurons and astrocytes were excluded for further analysis. Transcription factor–target regulatory network analysis was performed using the SCENIC method (Aibar et al., 2017). A genome search space between 500 bp and 10kb around the transcriptions start site was used.

## MERFISH

P35 CD1 mice were collected.10 µm tissue sections were obtained from along the rostro-caudal axis and adhered to the MERSCOPE Slide. The samples were permeabilized in 70% ethanol overnight, and followed by cell boundary staining. A 500-gene panel was generated based on enriched genes in each astrocyte and neuron cluster, with approximately 100 astrocyte genes and 400 neuron genes. The samples were then hybridized using the gene panel mix, followed by gel embedding and clearing steps. Images were processed using the MERSCOPE instrument and analysis computer, along with MERSCOPE visualizer software to streamline the acquisition of high-quality MERFISH data. Cells with a volume greater than 2000 µm^3, those with fewer than 100 transcripts, and those with DAPI scores less than 500 were excluded from the analysis. A total of 160,301 cells were used for downstream analysis, including 31,797 astrocytes.

### *In situ* hybridization

FPA-fixed brain sections were collected and preserved in a freezing buffer (250 ml buffer: 70g sucrose, 75 ml ethylene glycol, fill to 250 with 0.1M sodium phosphate buffer). These brain sections were pretreated using the RNAscope fluorescent multiplex assay which involved using RNA probes specific to *Slc6a11* and *Slc1a2*. For cell transplantation experiments, a multiplex RNAscope approach was employed by combining *Slc1a2* and *Slc6a11* probes with tdTomato immunostaining to confirm the identity of the labeled cells as transplanted cells.

### Cell transplantation

P3-P5 CD1 mice were used as recipients and Zic and Nkx2.1Cre;Ai14 mice were used as donors. Brains were sliced by using a 0.5 mm brain matrix, MS and LS regions were dissected in L15 medium (Gibco). Digest cells into single-cell suspension using papain. Dissociated cells were kept in ice cold L15 medium containing DNAse I (180μg/ml). Cells were then concentrated by spinning in a table centrifuge for five minutes at 800g (rcf), followed by the removal of the supernatant. The final cell pellet was resuspended and mixed in a final volume of 1-6μL of Leibovitz L-15 medium (L15). This concentrated cell suspension was loaded into beveled glass micropipettes (∼60-100 μm diameter, Wiretrol 5 μL, Drummond Scientific Company) prefilled with mineral oil and mounted on a microinjector as previously described (Wichterle, Garcia-Verdugo, Herrera, & Alvarez-Buylla, 1999). The viability and concentration of the cells in the glass micropipette were determined using a hemocytometer, loaded with 100 nl of cells diluted 200X in 10 μL of L15 medium and 10 μL trypan blue. Cells were injected into P4-P5 wild-type (CD1) mice. Prior to the injection of cells, the recipient mice were anesthetized by hypothermia (∼3-5 minutes) and positioned in a clay head mold to stabilizes the skull (Merkle, Mirzadeh, & Alvarez-Buylla, 2007). Micropipettes were positioned at an angle of 0 degrees from vertical in a stereotactic injection apparatus, and injected into the septum. The distance between the two eye corners was used to find the midline of the mouse brain. The injection coordinates to target the medial septum were 0 mm M/L from the midline, 2.75 mm A/P from the eye corner, and 3.2 mm D/V from the surface of the head. The coordinates to target the lateral septum were 0.5 mm M/L from the midline, 2.6 mm A/P from the eye corner, and 3.0 mm D/V from the surface of the head. After the injections were completed, the transplanted mice were placed on a warming pad to recover from hypothermia and return to their mothers afterwards. Brains were collected at P30. For morphology analysis, Z stack images were collected, maximizing the branch length of astrocytes using maximum intensity projection in ImageJ.

### Analysis of colocalized synaptic puncta

P21 GlastCreER;Ai14 mice were collected and sections were stained with antibodies against presynaptic and postsynaptic makers: VGluT1 and PSD95, or VGAT and Gephyrin. High magnification (100 × 1.44 NA objective plus 1.5X optical zoom) Z-stack images were obtained with a Leica SP8 confocal microscope. Each image includes individual astrocytes with 5 µm-thick Z stacks. The number of co-localized synaptic puncta on tdTomato+ astrocyte territories was analyzed using MATLAB (Xie et al., 2022).

### Quantification and statistical analysis

#### Statistical tests

Cell counting was analyzed manually with ImageJ/Fiji cell counter (National Institute of Health, USA). Analysis of cell density was done in ImageJ by dividing the septum into its medial and lateral nuclei based on DAPI staining, and the LS was divided into three subdivisions from dorsal to ventral. The area size of the region of interest was measured and the cells in these regions were counted. Quantification of synapse puncta in astrocyte territories and quantification of RNA puncta per unit area were carried out automatically using a data processing pipeline (see above) in MATLAB R2019b. All statistical tests were performed in GraphPad Prism 9. Parametric tests were used for normally distributed datasets while non-parametric tests were applied to data not normally distributed or data with a small sample size. For comparisons of two groups, paired and unpaired two-tailed Student’s t-tests or two-tailed Mann-Whitney tests were used. For comparisons of more than three groups, one-way ANOVA tests followed by Tukey’s or Kruskal-Wallis tests were used. For each experiment, the number of replicates (n) is indicated in figure legends. Significance was determined using the corresponding statistics test indicated in figure legends and reported as either p-value or: *p < 0.05, **p < 0.01, ***p < 0.001, ****p < 0.0001. When the p-value was greater than 0.05, it was stated as non-significant (n.s.).

---

**Figure S1, related to Figure 1:**
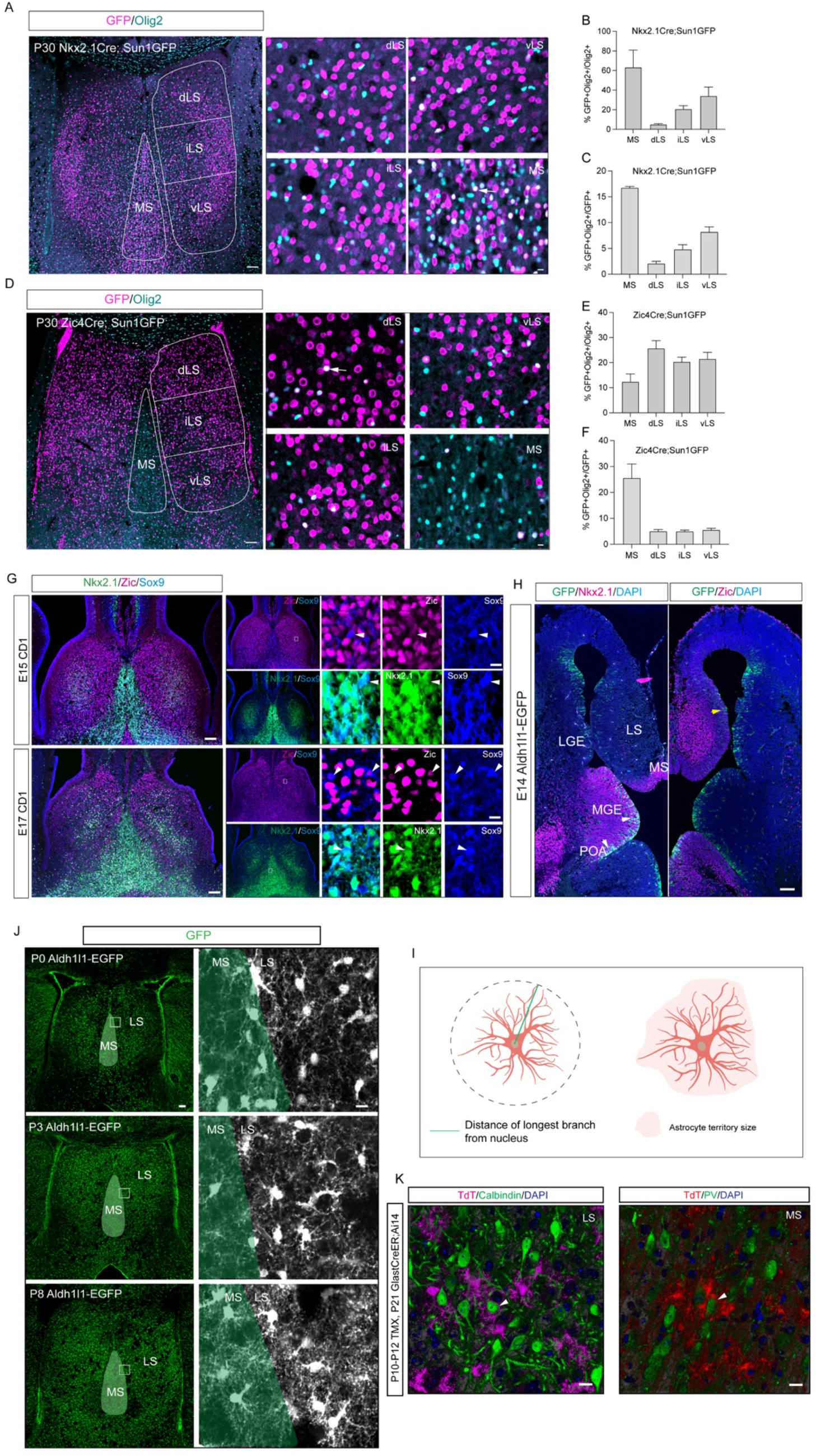
Nkx2.1 and Zic4 lineages-derived astrocytes occupy MS and LS respectively. **A**) Immunostaining for Olig2 (cyan) and GFP (magenta) in P30 Nkx2.1Cre; Sun1GFP septum. Subdivisions of the septum are indicated with white lines. White arrows indicate Nkx2.1-derived oligodendrocytes. Scale bar: left: 100 µm, right: 10 µm. **B**) Proportion of Nkx2.1-derived oligodendrocytes in total oligodendrocytes. N=5 mice. **C)** Proportion of Nkx2.1-derived oligodendrocytes in total Nkx2.1-derived cells. N=4 mice. **D**) Immunostaining for GFP (magenta) and Olig2 (cyan) in P30 Zic4Cre; Sun1GFP mice. Subdivisions of the septum are indicated with white lines. White arrows indicate cells with overlapping signals. Scale bar: left: 100 µm, right: 10 µm. **E**) Proportion of Zic4-derived oligodendrocytes in total oligodendrocytes. N=5 mice. **F)** Proportion of Zic4-derived oligodendrocytes in total Zic4-derived cells. N=5 mice. **G**) Immunostaining for Nkx2.1 (green), Zic (magenta), and Sox9 (blue) in coronal sections of E15 and E17 CD1 mice. White boxes indicate the selected regions for high magnification images (right panels. White arrows indicate the Zic+/Sox9+ and Nkx2.1+/Sox9+ cells. Scale bar: left: 100 µm, right: 10 µm. **H**) Immunostaining for GFP (green), Nkx2.1 (magenta), Zic (magenta), and DAPI (blue) in horizontal sections of E14 Aldh1l1-EGFP mice. Lateral ganglionic eminence (LGE); medial ganglionic eminence (MGE); preoptic area (POA). White arrows indicate the Nkx2.1+/GFP+ astrocyte progenitors and a yellow arrow indicates Zic+/GFP+ astrocyte progenitors. Scale bar: 100 µm. **I**) Schematic of the methods for analyzing astrocyte longest branch and territory size. **J**) Immunostaining for GFP in P0, P3, and P8 Aldh1l1-EGFP mice, MS and LS boundaries were distinguished by the density of DAPI. White boxes indicate the selected regions for high magnification images (right panel). Scale bar: left: 100 µm, right: 10 µm. **K**) Immunostaining for tdTomato (red/magenta), Calbindin (green)(left), PV (green)(right), and DAPI (blue) in tamoxifen-induced P21 GlastCreER; Ai14 mice (tamoxifen injected at P10-P12). White arrows indicate Calbindin+ and PV+ neurons tightly ensheathed by adjacent tdTomato+ astrocytes in the LS and MS respectively. Scale bar: 10 µm.

**Figure S2, related to Figure 2:**
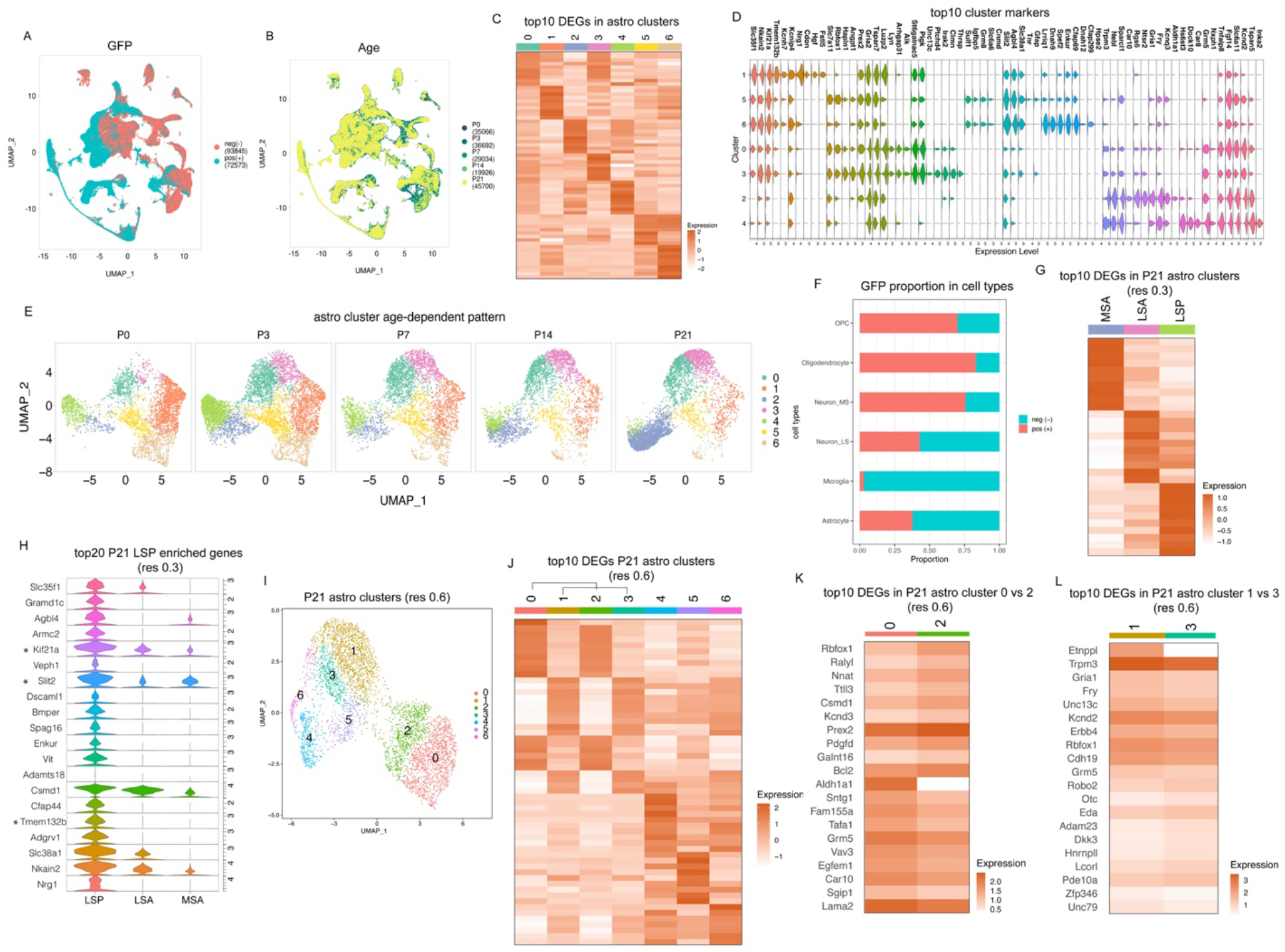
Septal astrocytes display lineage-specific molecular profiles. **A**) UMAP plot of Nkx2.1-derived (GFP+) and non-Nkx2.1-derived (GFP-) cells. **B**) UMAP analysis of total cells at different ages. **C)** Heatmap showing the top 10 DEGs in seven subgroups of astrocytes. **D**) Top 10 DEGs enriched in individual astrocyte clusters. **E**) The dynamic changes in subsets of astrocyte populations over ages. **F**) The proportion of GFP+ and GFP-cells in each cell type. **G**) Heatmap showing the top 10 DEGs in P21 astrocyte clusters at a 0.3 resolution level. **H**) A violin plot showing the top 20 genes highly enriched in the LSP cluster. * indicates genes highlighted in the text. **I**) UMAP plot showing 7 astrocyte subclusters in P21 at a 0.6 resolution level. **J**) Heatmap showing the top 10 DEGs in seven subgroups of P21 astrocyte at a 0.6 resolution level. **K**) Heatmap showing the top 10 DEGs in P21 astrocyte cluster 0 versus cluster 2 (corresponding to figure S2I) at a 0.6 resolution level. **L**) Heatmap showing the top 10 DEGs in P21 astrocyte cluster 1 versus cluster 3 (corresponding to figure S2I) at a 0.6 resolution level.

**Figure S3, related to Figure 2:**
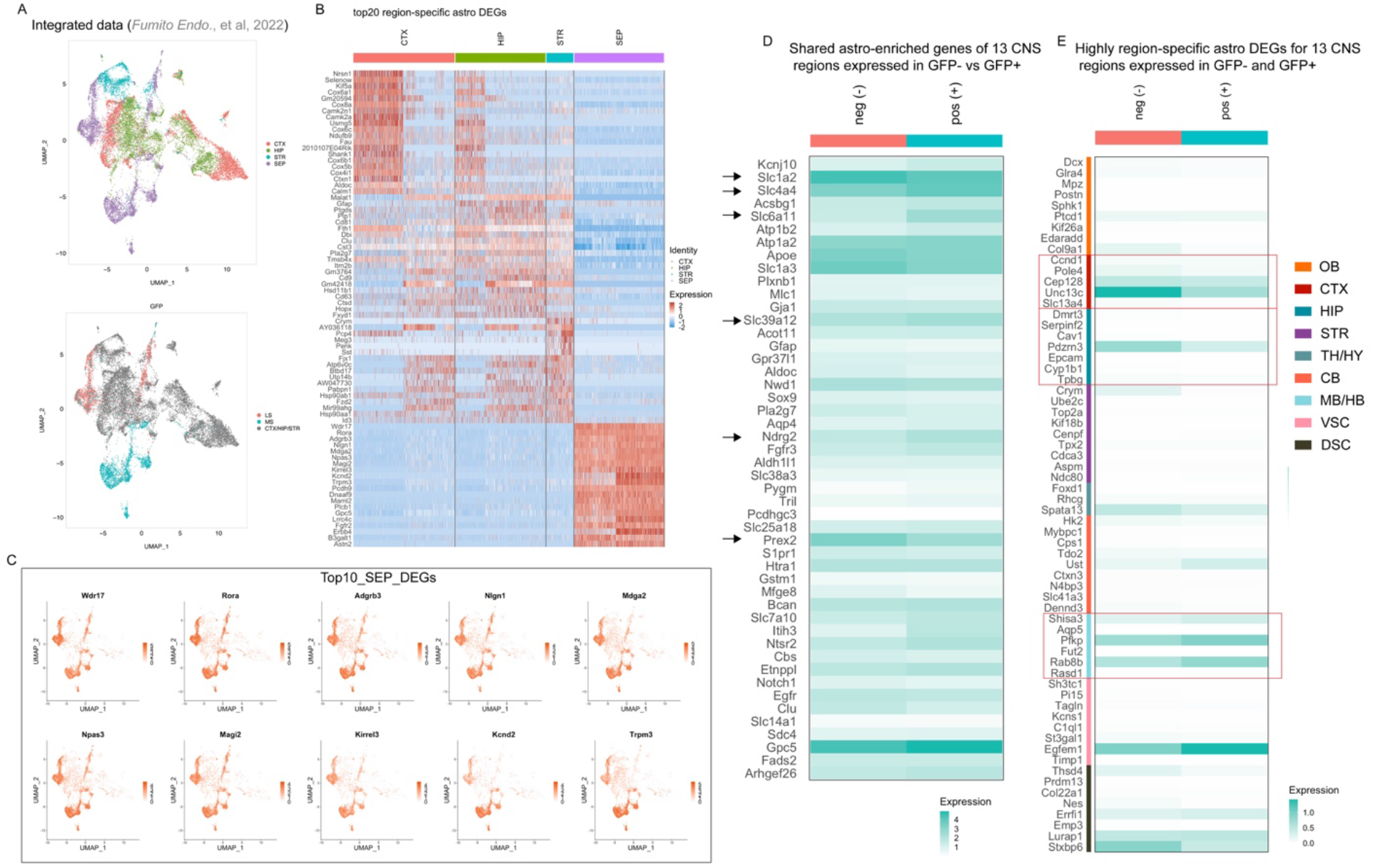
Septal astrocytes exhibit region-specific molecular signatures. **A**) UMAP plot showing astrocyte clusters in four distinct regions (top) and segregation between MS astrocytes and other regions based on GFP expression (bottom). CTX = cortex, HIP = hippocampus, STR = striatum, SEP = septum. **B**) Heatmap showing the top 20 region-specific enriched genes in the CTX, HIP, STR and SEP. **C**) Feature plots of the top 10 septal DEGs. **D**) Heatmap showing genes shared across 13 CNS regions (the olfactory bulb (OB), motor cortex (MCX), somatosensory cortex (SCX), visual cortex (VCX), hippocampus (HIP), striatum (STR), thalamus (TH), hypothalamus (HY), cerebellum (CB), midbrain (MB), hindbrain (HB), ventral spinal cord (VSC) and dorsal spinal cord (DSC)) expressed in P21 GFP+ and GFP-septal astrocytes. Arrows indicate highly differentially expressed genes. **E**) Heatmap plot showing the enrichment of region-specific DEGs from 13 CNS regions (indicated in figure legend S3D) in GFP- and GFP+ septal astrocytes. Red boxes indicate genes enriched in CTX, HIP, and MB/HB regions.

**Figure S4, related to Figure 2:**
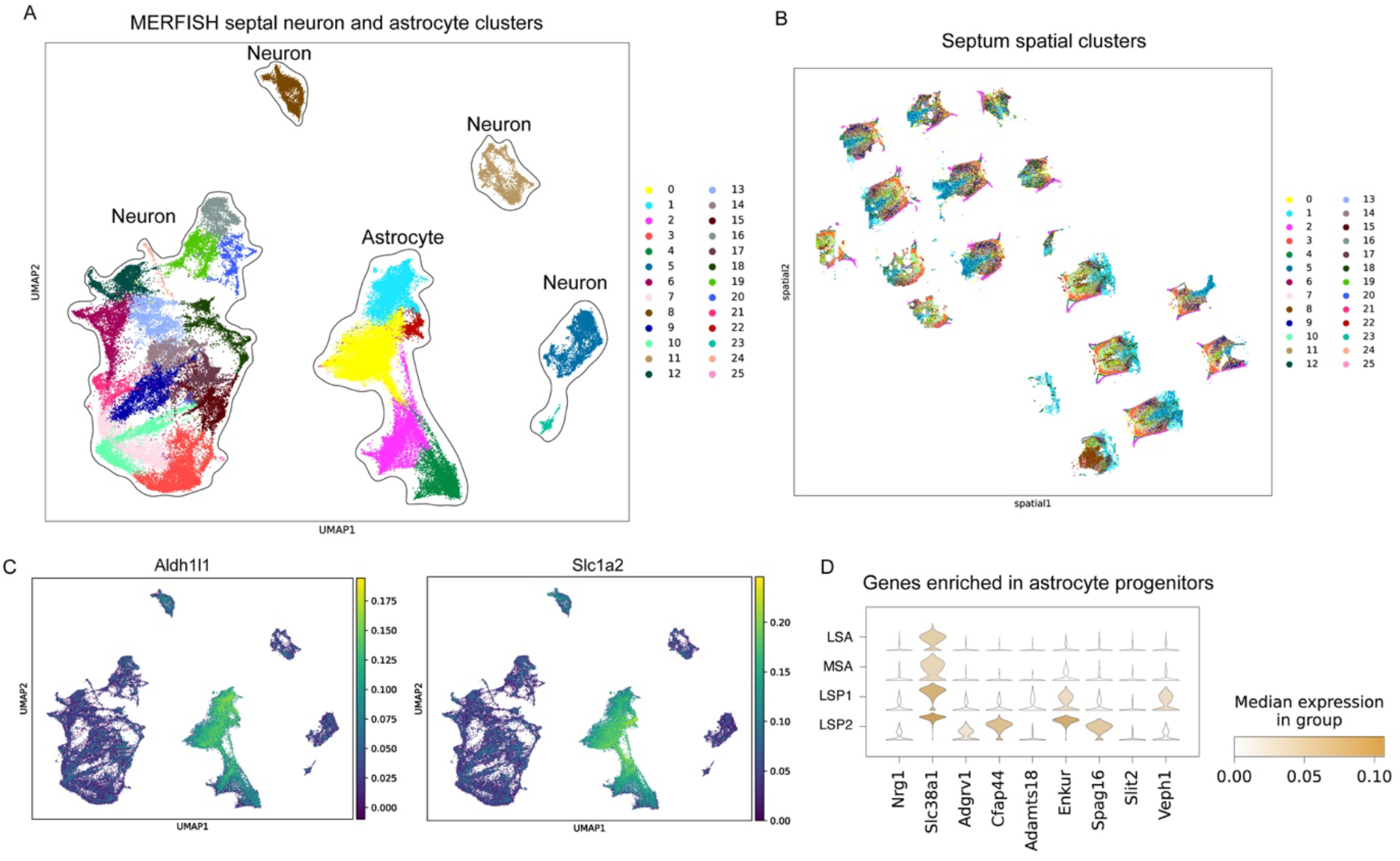
MERFISH analysis of P35 septal cells. **A**) UMAP plot of MERFISH data revealing 25 subclusters of P35 septal astrocytes and neurons. **B**) Spatial transcriptomics analysis showing 25 subcluster locations in the septum through the rostro-caudal axis. **C**) Astrocyte clusters in MERFISH were identified by general astrocyte markers: *Aldh1l1* and *Slc1a2*. **D**) Genes enriched in LSP cluster (snRNAseq, figure S2H) show high enrichments in MERFISH clusters: LSP1 and LSP2.

**Figure S5, related to Figure 3:**
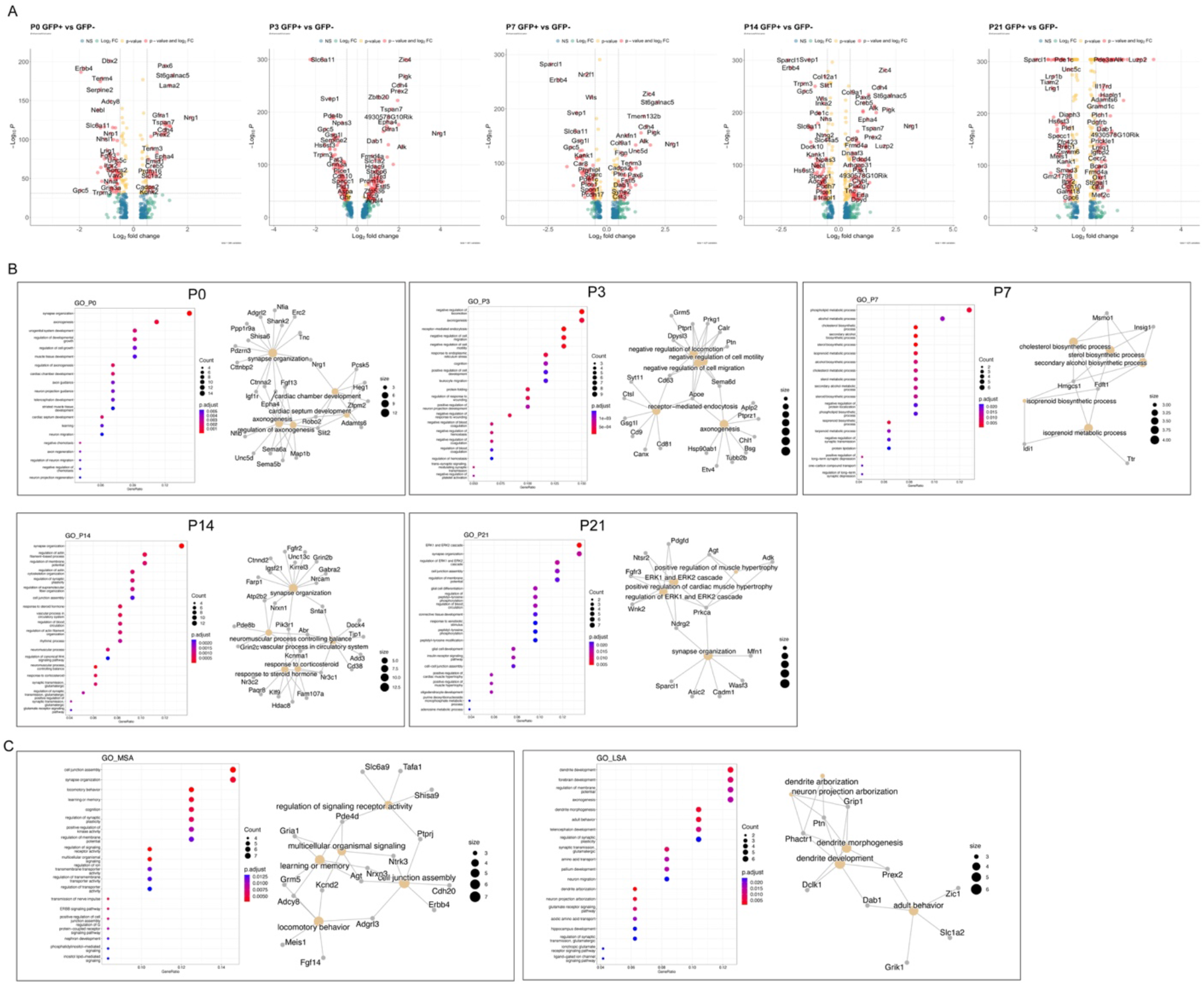
Age-dependent molecular profiles and function in septal astrocytes. **A**) Volcano plots showing the genes enriched in GFP+ versus GFP-developing astrocytes. **B**) Gene ontology analysis (top 20 gene terms) of DEGs in septal astrocytes at different ages. **C**) Gene ontology analysis (top 20 gene terms) of DEGs in P21 MS and LS astrocytes.

**Figure S6, related to Figure 3:**
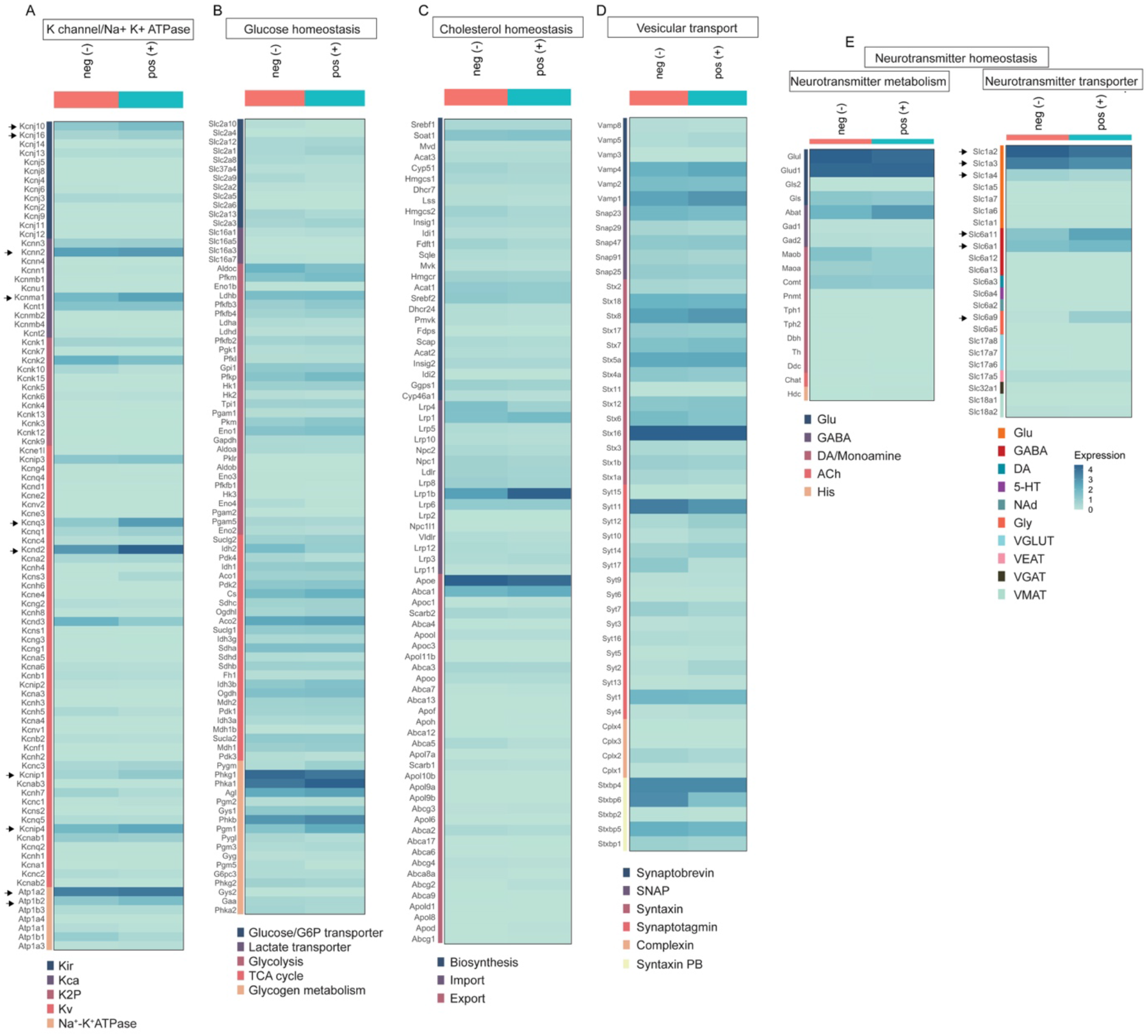
Expression of genes regulating astrocytic core function in septal astrocytes. **A-E**) Heatmaps showing the average expression of genes for potassium channels and Na+-K+ ATPases (**A**), glucose homeostasis (**B**), cholesterol homeostasis (**C**), vesicular transport (**D**), and neurotransmitter homeostasis (**E**) in P21 GFP+ vs GFP-astrocytes. Kir: inwardly rectifying K+ channel, Kca: calcium-activated K+ channel, K2P: two-pore domain K+ channel, Kv: voltage-gated K+ channel, Glu: glutamate, DA: dopamine, Ach: acetylcholine, 5-HT: serotonin, His: histamine, Ad: Adrenaline, Gly: glycine, VGLUT: glutamate vesicular transporter, VEAT: vesicular excitatory amino acid transporter, VGAT: vesicular GABA transporter, VMAT: vesicular monoamine transporter, VAChT: vesicular acetylcholine transporter. Black arrows indicate the highly enriched genes in either GFP- or GFP+ astrocytes.

**Figure S7, related to Figure 3:**
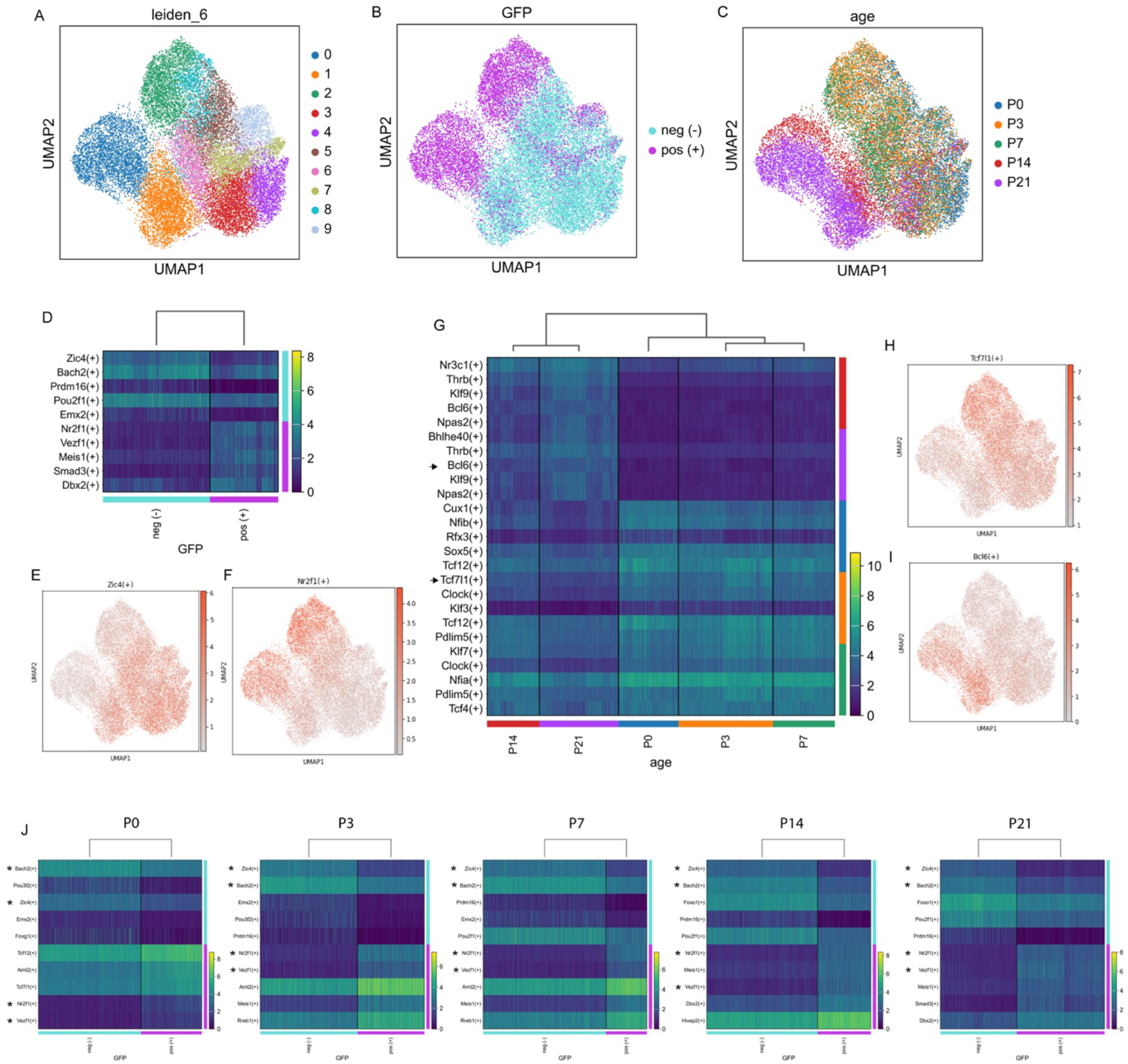
SCENIC analysis for regulon networks in septal astrocytes. **A**) UMAP plot defines 10 subclusters of astrocytes regarding distinct regulon networks. **B**) GFP+ and GFP-astrocytes are distinguished from each other based on regulon networks. **C**) Age-dependent regulon networks in septal astrocytes through development. **D**) Top 5 regulons enriched in MS and LS astrocytes. **E, F**) Feature plots showing Zic4 (**E**) and Nr2f1 (**F**) regulons highly enriched in LS and MS astrocytes respectively. **G**) The top enriched age-dependent regulons in septal astrocytes. Arrows indicate the highlighted genes in the text. **H, I**) Tcf7l1 (**H**) and Bcl6 (**I**) are enriched in septal astrocytes at early and late ages. **J**) Top 5 enriched regulons in GFP+ and GFP-astrocytes at each age. * indicates the conserved regulons at different ages.

**Figure S8, related to Figure 4:**
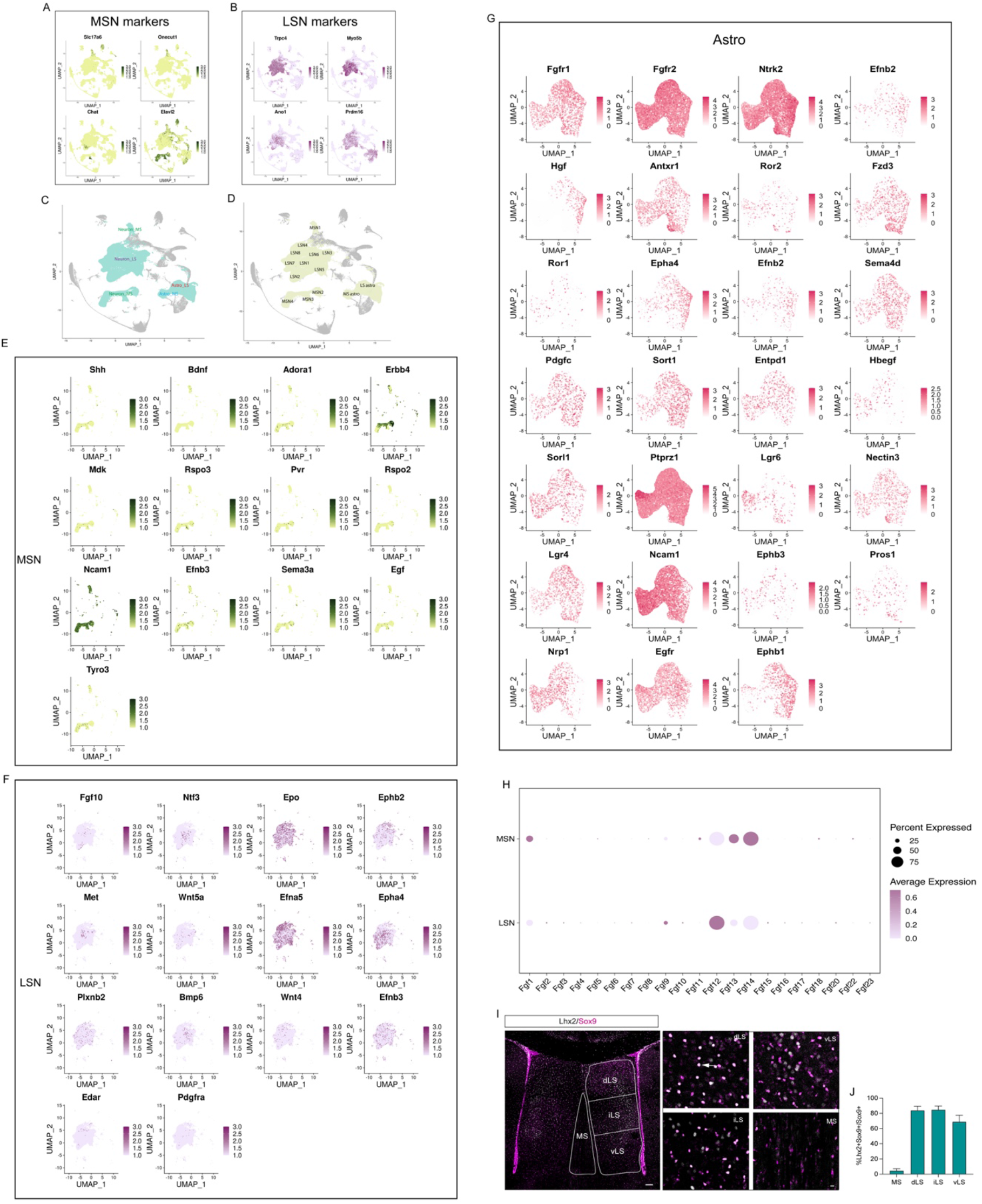
Ligand-receptor expression patterns in the septum. **A, B**) Representative genes define MS (**A**) and LS (**B**) neurons. MSN: medial septal neuron; LSN: lateral septal neuron. **C**, **D**) Highlighted neuron and astrocyte subclusters are processed by CellphoneDB analysis. **E-G**) Expression patterns of ligands and receptors in MSN (**E**), LSN (**F**), and septal astrocytes (Astro) (**G**). **H**) Expression patterns of Fgf family members in septal neurons. **I**) Immunostaining for Lhx2 (grey) and Sox9 (magenta) in P30 wild-type septum. Subdivisions of the septum are indicated with white lines. White arrows indicate cells with overlapping signals. Scale bars: left, 100 µm; right, 10 µm. **J**) Proportion of Lhx2+ astrocytes in total astrocytes across different regions. N=5 mice.

**Figure S9, related to Figure 5:**
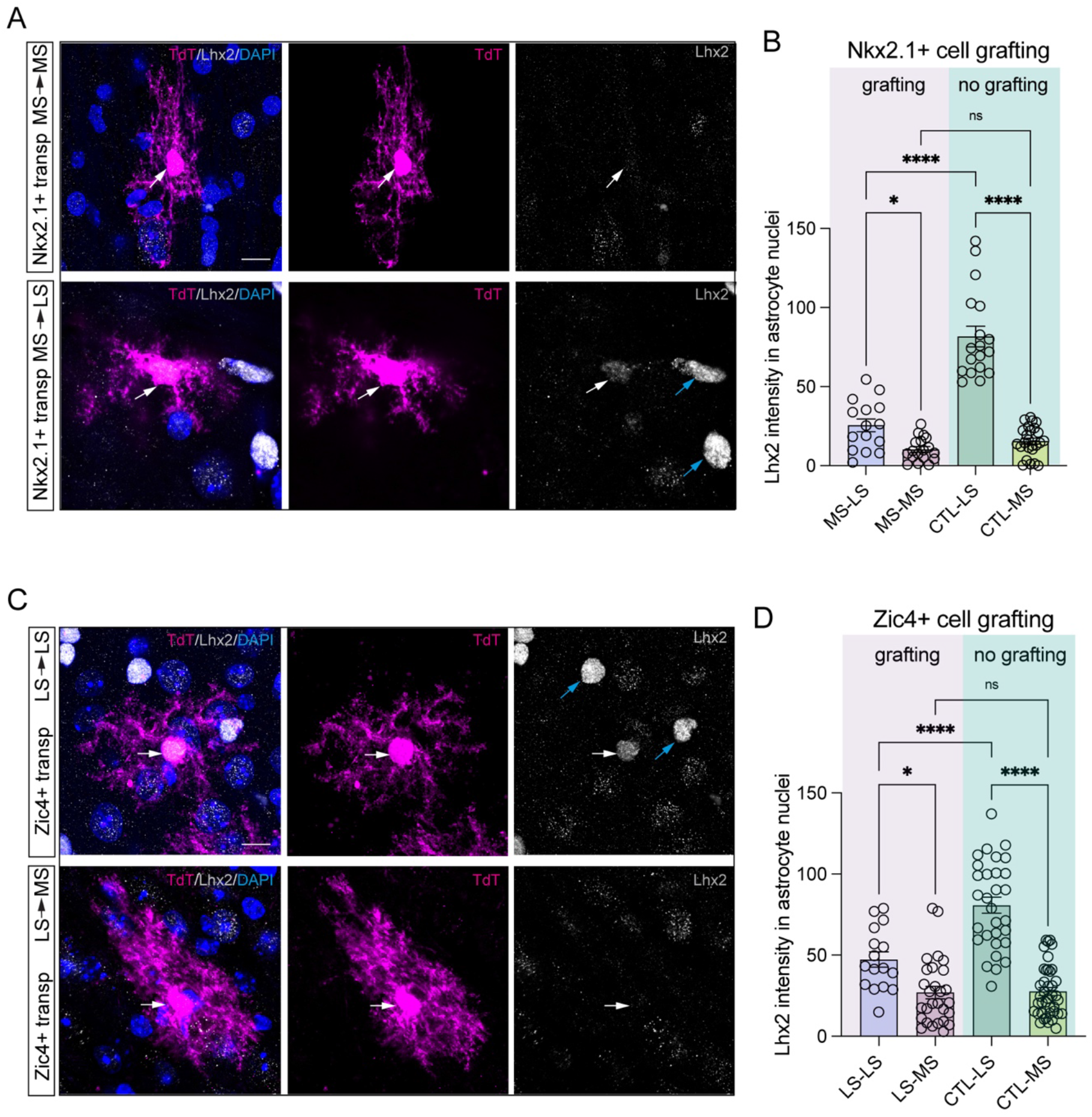
Grafted astrocytes adopt local astrocyte molecular features. **A**) Immunostaining for Lhx2 (grey), tdTomato (magenta) and DAPI (blue) in transplanted Nkx2.1-derived astrocytes. White arrows indicate transplanted astrocytes; blue arrows indicate host astrocytes. **B**) Quantification of Lhx2 intensity on astrocytes nuclei on both grafting and no grafting groups. N=4-6 transplants, 15-25 astrocytes were analyzed for each group. **C**) Immunostaining for Lhx2 (grey), tdTomato (magenta) and DAPI (blue) in transplanted Zic4-derived astrocytes. White arrows indicate transplanted astrocytes and blue arrows indicate host astrocytes. **D**) Quantification of Lhx2 intensity on astrocyte nuclei on both grafting and no grafting groups. N=4-6 transplants, 16-40 astrocytes were analyzed for each group.

**Table S1, related to Figure 3: SCENIC analysis revealing the transcription factor– target regulatory networks in septal astrocytes.** “Genes” indicate the list of target genes of each transcription factor. “Weights” indicate the close linkage between each target and the corresponding transcription factor.

